# *Methanobrevibacter smithii* associates with colorectal cancer through trophic control of the cancer bacteriome

**DOI:** 10.1101/2025.06.12.659283

**Authors:** Rokhsareh Mohammadzadeh, Alexander Mahnert, Tamara Zurabischvili, Lisa Wink, Christina Kumpitsch, Hansjoerg Habisch, Jannik Sprengel, Klara Filek, Polona Mertelj, Dominique Pernitsch, Kerstin Hingerl, Gregor Gorkiewicz, Christian Diener, Alexander Loy, Dagmar Kolb, Christoph Trautwein, Tobias Madl, Christine Moissl-Eichinger

## Abstract

The human gut is colonized by trillions of microbes that influence the health of their human host. Whereas many bacterial species have now been linked to a variety of different diseases, the involvement of Archaea in human disease remains elusive. Here we searched for gut archaeal signatures of disease by screening 19 cross-sectional clinical studies comprising more than 1,800 individuals. We found that associations between Archaea and medical disorders are common but highly variable and are dominated by a significant increase of *Methanobrevibacter smithii* in colorectal cancer (CRC) patients. Metabolic modelling and *in vitro* co-culture identified distinct mutualistic interactions of *M. smithii* with CRC-causing bacteria such as *Fusobacterium nucleatum*, including metabolic enhancement. Metabolomics further revealed archaeal-derived compounds with tumor-modulating properties. This provides the first mechanistic link between human gut archaeome and CRC and highlights its role in modulating health in humans through trophic control of the resident bacteriome.

## Introduction

Alterations in the gastrointestinal microbiome have been implicated in a wide range of diseases, including inflammatory bowel diseases (IBD), metabolic disorders like obesity and type 2 diabetes (T2D), cardiovascular conditions, and neurodegenerative or neuropsychiatric diseases including Parkinson’s disease (PD), Alzheimer’s disease (AD), schizophrenia (SCZ), and multiple sclerosis ^1–8^. Establishing mechanistic links between specific microbes and disease phenotypes remains a major challenge due to the inherent complexity and interdependence of microbial ecosystems. Microbiomes are shaped by intricate networks of cross-feeding, competition and host interactions across domains of life, including a complex interplay of bacteria, fungi, viruses – and archaea.

Archaea, in particular methane-forming representatives, can constitute up to 4% of the gut microbiome^9^, but remain severely under-researched. The most abundant genus is *Methanobrevibacter,* whose representatives are generally considered commensal and have not been linked to pathogenicity in humans or other hosts^10^. Yet, their metabolic activity - primarily the consumption of bacterial fermentation products (H₂, CO₂) to produce methane - positions them as key microbial interactors^11^. *Methanobrevibacter* abundance has been associated with beneficial outcomes, including higher short-chain fatty acid levels, reduced body mass index (BMI), and increased longevity^9,12–14^ Archaeal depletion has been observed in several disorders such as IBD^15^, obesity^16^, and irritable bowel syndrome with diarrhea (IBS-D)^17,18^. Conversely, elevated *Methanobrevibacter* levels have also been linked to constipation-dominant IBS (IBS-C)^19^, intestinal methanogen overgrowth (IMO, a small intestinal bacterial overgrowth (SIBO) subtype)^20,21^, and in some reports, colorectal cancer (CRC)^22^. These associations suggest that methanogens may modulate host physiology both directly, through metabolite production, and indirectly, by shaping bacterial activity through otherwise inhibiting H_2_ removal.

Despite their potential significance, archaea are largely neglected in most microbiome studies due to methodological limitations. Standard 16S rRNA gene-targeted primers lack archaeal coverage, reference genome databases remain incomplete, cultivation is tricky, and many computational pipelines are not optimized to detect archaeal signatures^23,24^. Most existing data on archaea come from using low-resolution methods, such as breath methane testing or 16S rRNA gene profiling, which cannot resolve species-level taxa or infer metabolic functions. As a result, key hypotheses remain untested: that distinct diseases may induce characteristic shifts in gut archaeome (the archaeal community residing in the human gut); that methanogens may be selectively enriched or depleted depending on the disease context; that certain archaeal taxa could serve as reliable, disease-specific biomarkers across diverse cohorts; and that archaeal metabolites - whether unique to this domain or produced via syntrophic interactions - may exert biologically relevant effects on host physiology and disease processes.

To address these knowledge gaps, we performed a multi-level study, starting with the first comprehensive meta-analysis of the human gut archaeome across multiple diseases, leveraging high-quality shotgun metagenomes from 1,882 fecal samples derived from 19 studies spanning 12 countries. This dataset covered a spectrum of conditions: CRC, T2D, CD, UC, MS, AD, SCZ, and PD. Using a unified analytical framework, we systematically quantified archaeal prevalence, identified disease-specific patterns, and assessed the significance of these associations across cohorts while adjusting for major confounders including age, sex, and BMI. Observed correlations with CRC were further addressed by genome-scale metabolic modeling and archaeal-bacterial co-culture experiments, involving CRC-associated pathogens such as *Fusobacterium nucleatum*. Functional interactions were resolved by metabolomics, revealing that *M. smithii* engages in complex exchanges that have potential to shape the tumor microenvironment.

Our findings position *M. smithii* as an active metabolic and ecological contributor within CRC-associated microbial networks. By redefining its role from a passive H_2_ scavenger to a dynamic modulator of microbial interactions and host-relevant metabolites, this study highlights the importance of integrating archaeal functions into microbiome-disease frameworks.

## Materials and Methods

### Dataset collection

Relevant metagenomic investigations were identified through targeted keyword searches for diseases previously linked to gut methanogens^1–8,25^, specifically colorectal cancer (CRC), inflammatory bowel disease (IBD), type 2 diabetes (T2D), multiple sclerosis (MS), Alzheimer’s disease (AD), schizophrenia (SCZ), and Parkinson’s disease (PD) (Table 1). A systematic PubMed search was performed for English-language observational studies and meta-analyses published from 2000 until August 30, 2024. The search query used was: ("Disease of Interest") AND ((gut AND metagenome AND microbiome) OR fecal OR shotgun OR microbiota OR "whole genome"), with “Disease of Interest” encompassing CRC, IBD, PD, AD, T2D, MS, and SCZ. Additionally, reference lists from the identified articles and related meta-analyses were screened to identify additional studies. Additionally, the NCBI BioProjects database was screened for sequencing datasets using disease-specific terms (e.g., “CRC gut”), with filters applied for the “metagenome” data type and the human organism.

**Table 1.**
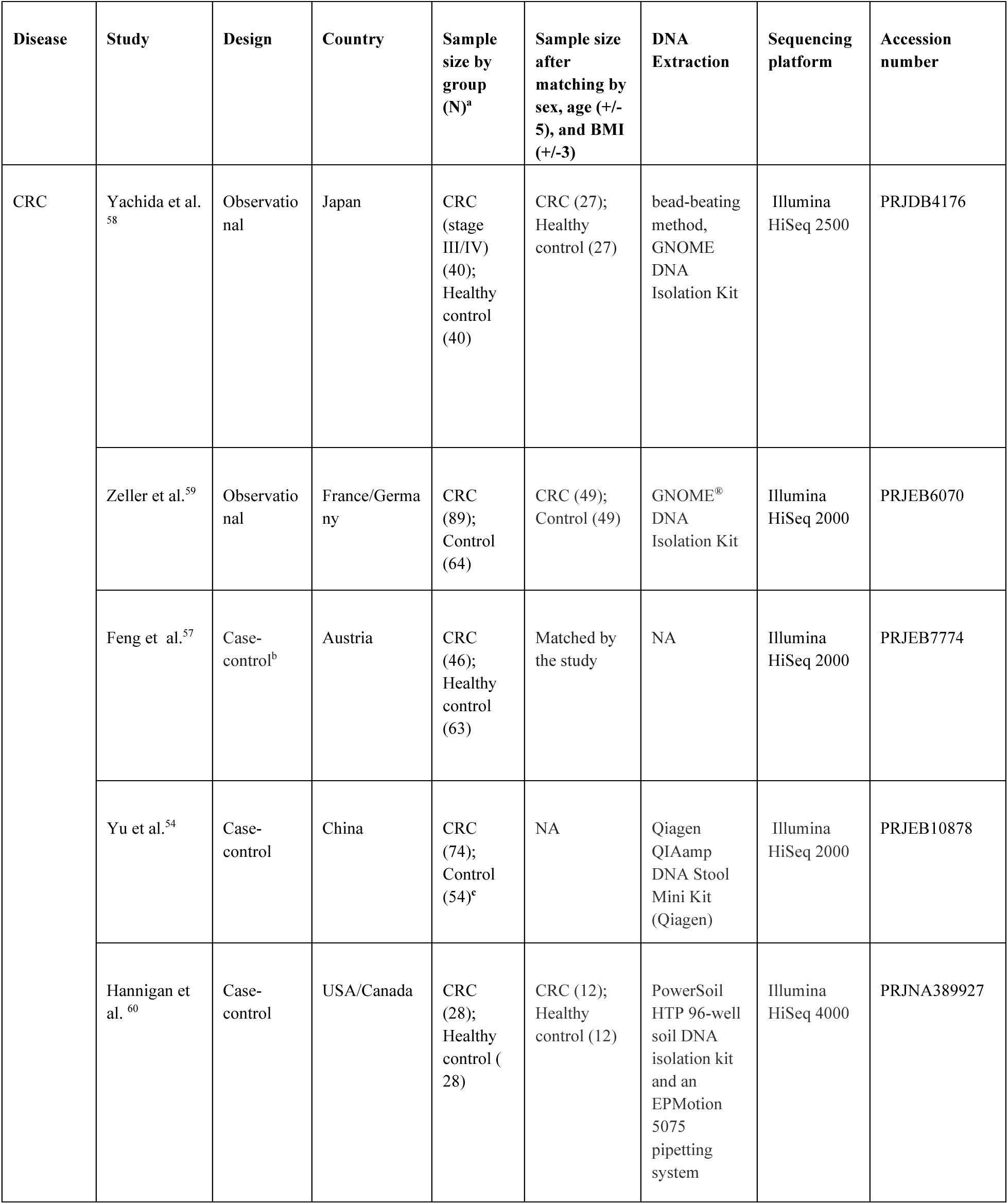
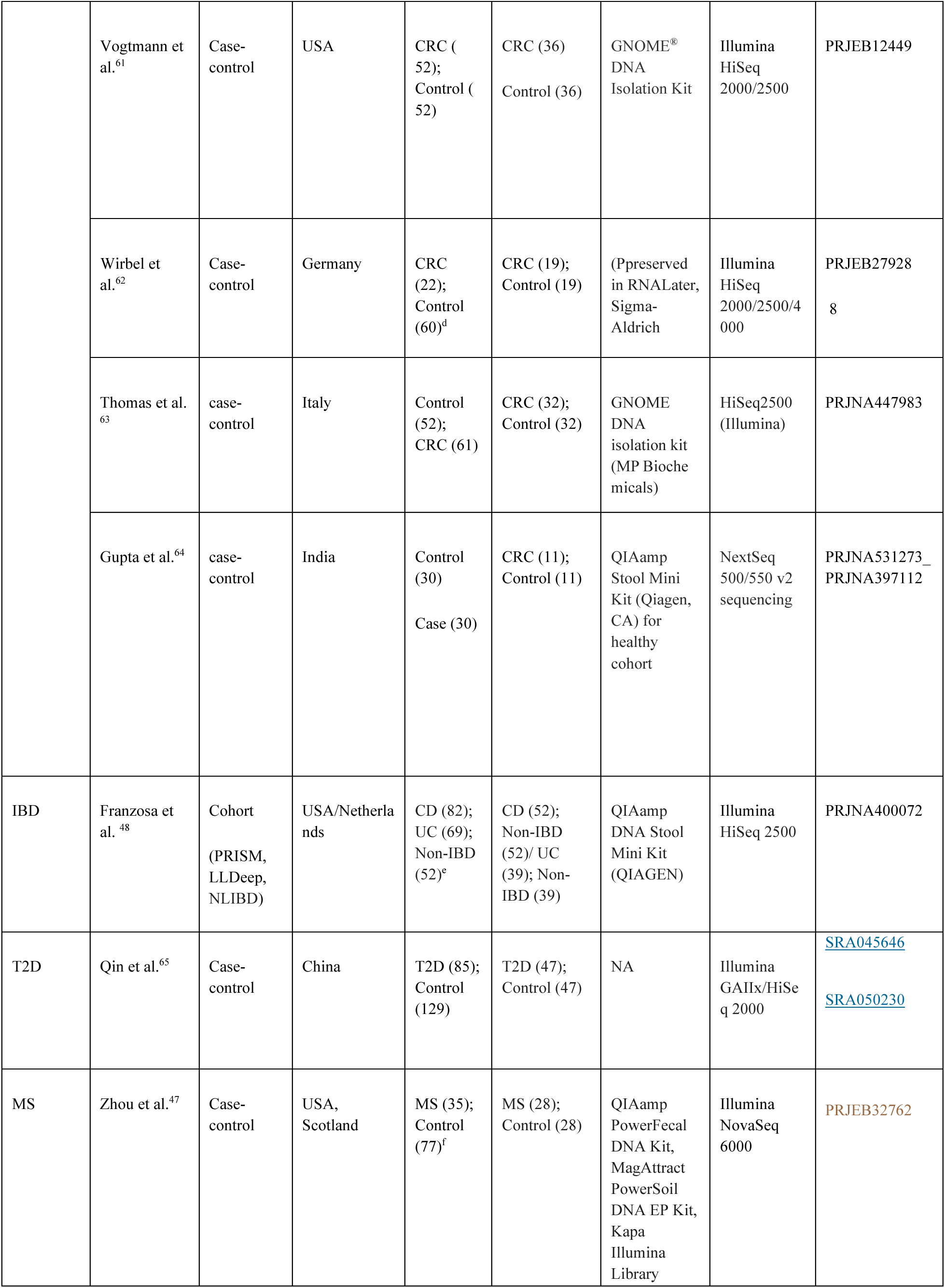
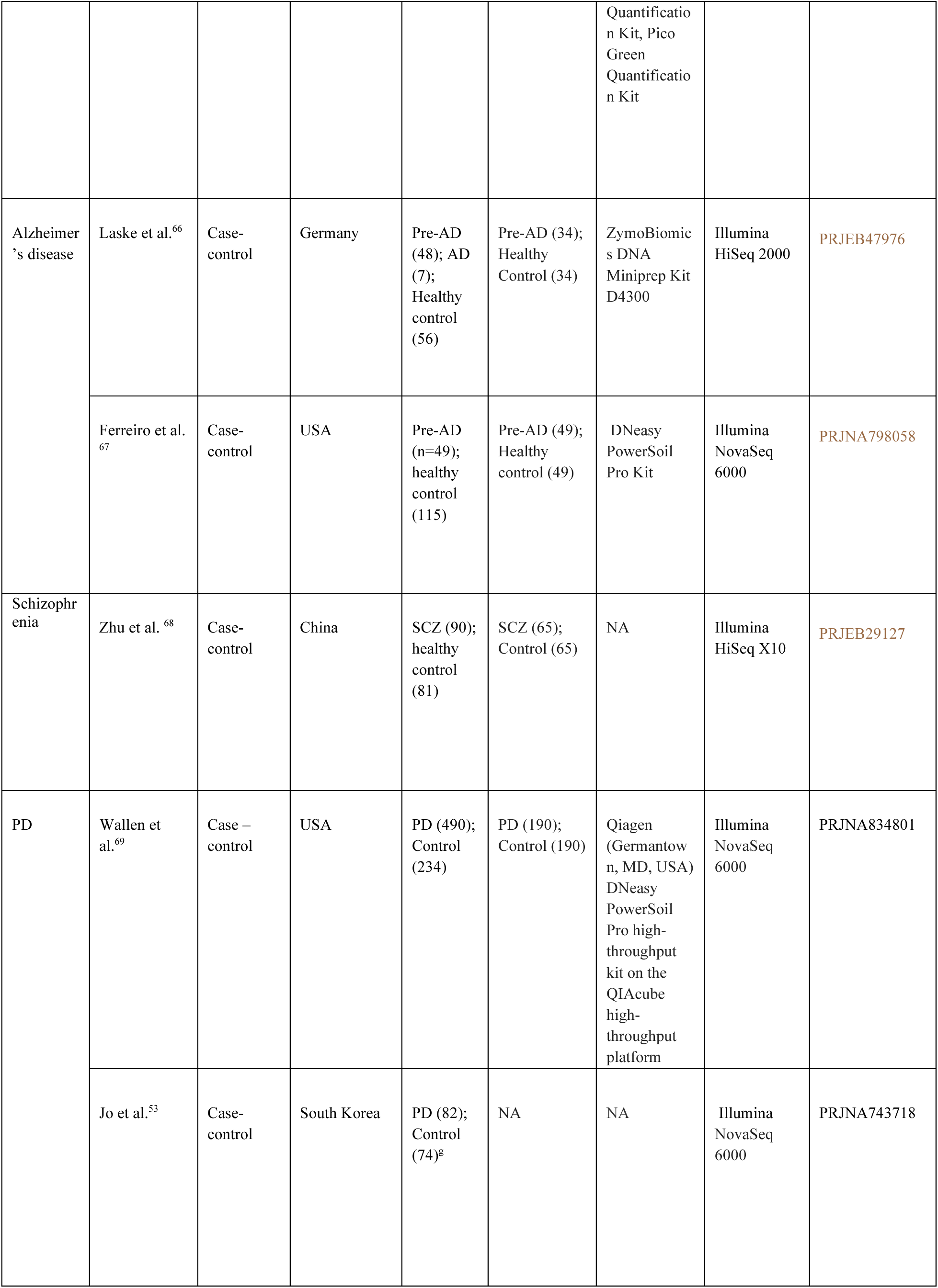
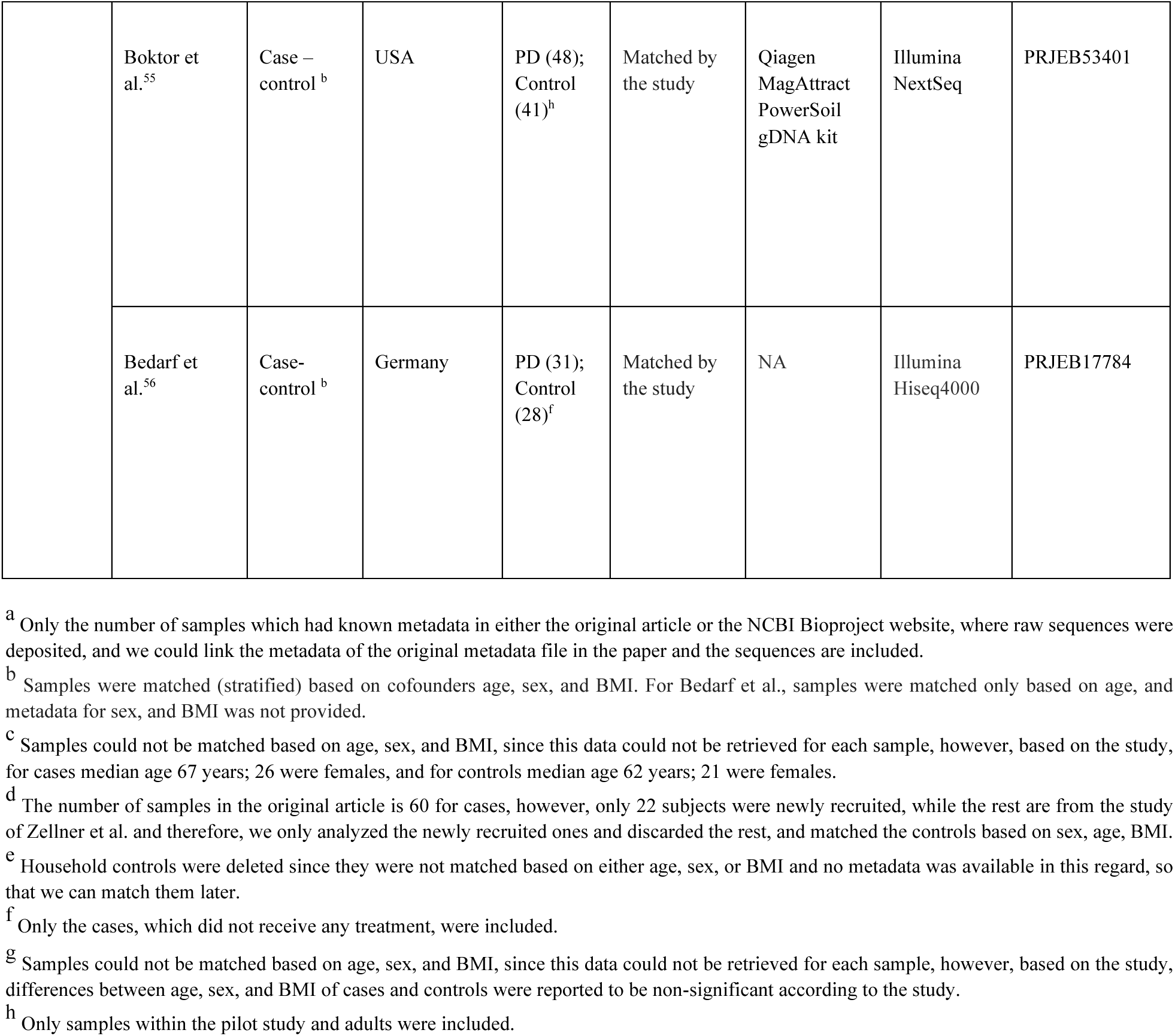
Details of the studies and fecal metagenome samples included in the meta-analysis.

Studies employing shotgun metagenomic sequencing were considered. Only fecal shotgun metagenomic datasets were included, while studies with fewer than 20 case subjects or those involving participants under 18 years were excluded. Case subjects were defined as individuals with explicit disease diagnoses provided in the study metadata or NCBI BioProject records, whereas control subjects were those clearly described as healthy or designated as controls. Controls were defined as samples from individuals without a positive disease diagnosis. Thus, it should be noted that controls do not necessarily present healthy individuals, but rather individuals without a specific diagnosis. Furthermore, studies were required to report metadata on sex, age, and/or BMI, or to have used matched cohorts on at least two of these covariates. Studies that would have required additional ethical approvals or had restricted data access were omitted.

For CRC investigations, only samples from patients with confirmed CRC diagnoses were incorporated, excluding those with small or large adenomas. In the case of MS, inclusion was limited to two cohorts (from San Francisco, USA, and Edinburgh, Scotland) that consisted of treatment-naïve individuals and met the required sample size threshold, since treatment in these patients affects the abundance of the archaeome^26^. For AD, only pre-AD samples were considered given the limited number of diagnosed cases. Sequence data were further limited to paired-end read libraries, thereby excluding datasets that combined single-end and paired-end reads. When available, sample covariates were extracted from the original study metadata, and for individuals with multiple SRA accessions, only the first available metagenome sample was used. Finally, any samples lacking accessible metadata from either the NCBI BioProject records or the published article were excluded.

Following these criteria, a total of 2,881 samples from 19 studies across 12 countries were identified (Table 1). Further details on study selection and the number of studies considered are provided in Figure 1 and Figure S1.

**Figure 1.**
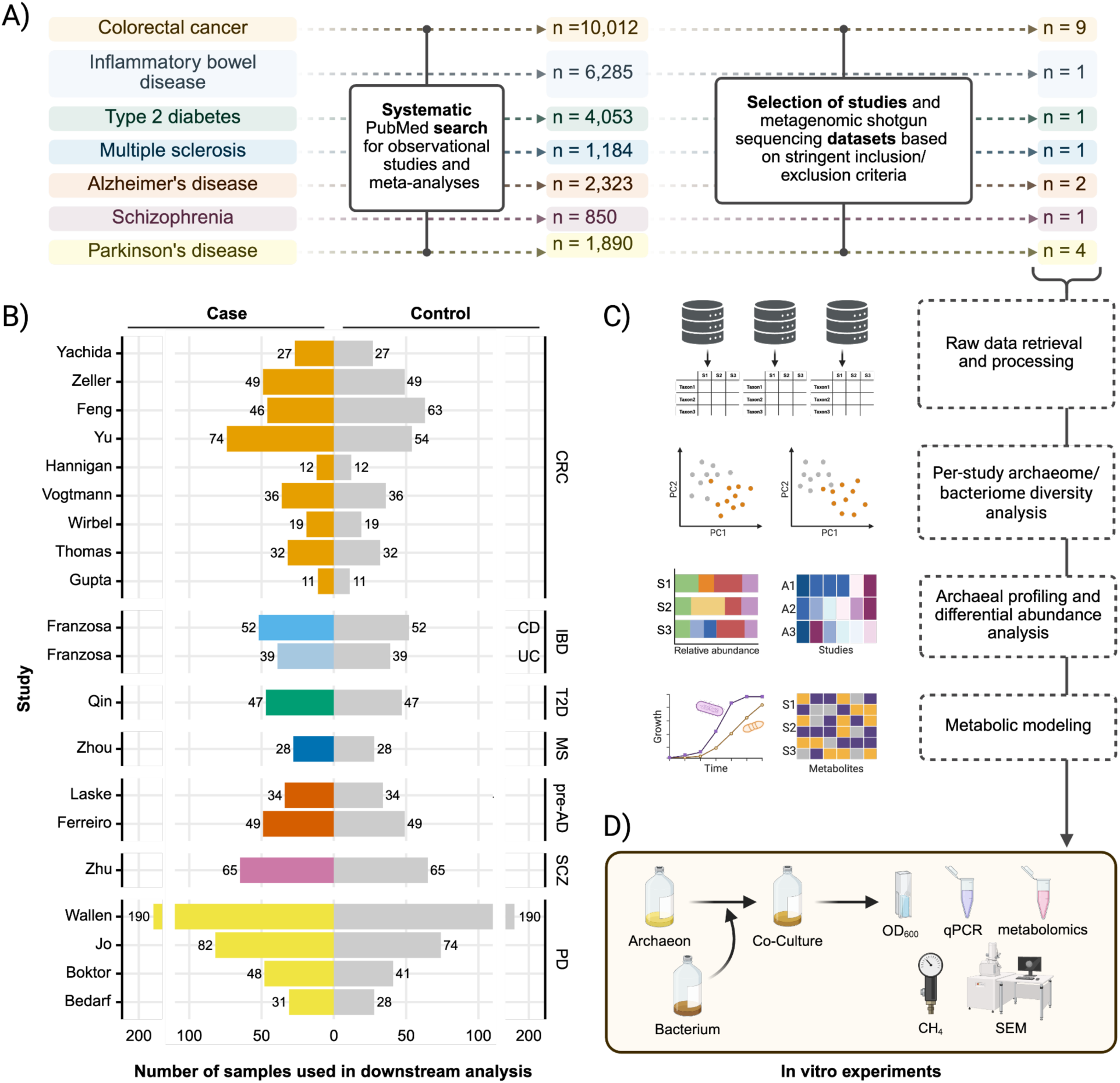
Schematic overview of the study. A) Metagenomic sequencing datasets were selected using stringent inclusion and exclusion criteria. B) Nine studies were retained for downstream analysis, and samples were adjusted for confounders such as sex, age, and BMI when possible. C) Raw reads were retrieved, quality-controlled, and filtered to remove human sequences, followed by taxonomic classification using Kraken2 and Bracken. Archaeal profiles were then extracted, and differential abundance testing was performed. Metabolic modeling was used to explore metabolite exchange between *Methanobrevibacter smithii* and CRC-associated bacteria. D) *In vitro* experiments were conducted to understand the dynamics between *M. smithii* and CRC-associated bacteria. CRC, Colorectal cancer; CD, Crohn’s disease; UC, Ulcerative Colitis; T2D, Type 2 Diabetes; MS, Multiple Sclerosis; pre-AD, pre-Alzheimer’s Disease; SCZ, Schizophrenia; PD, Parkinson’s Disease. Figure panels A, C, and D were created using BioRender. Panel B visualization was performed using the ggplot2 package, and axis breaks were introduced using ggbreak^70,71^.

**Figure S1.**
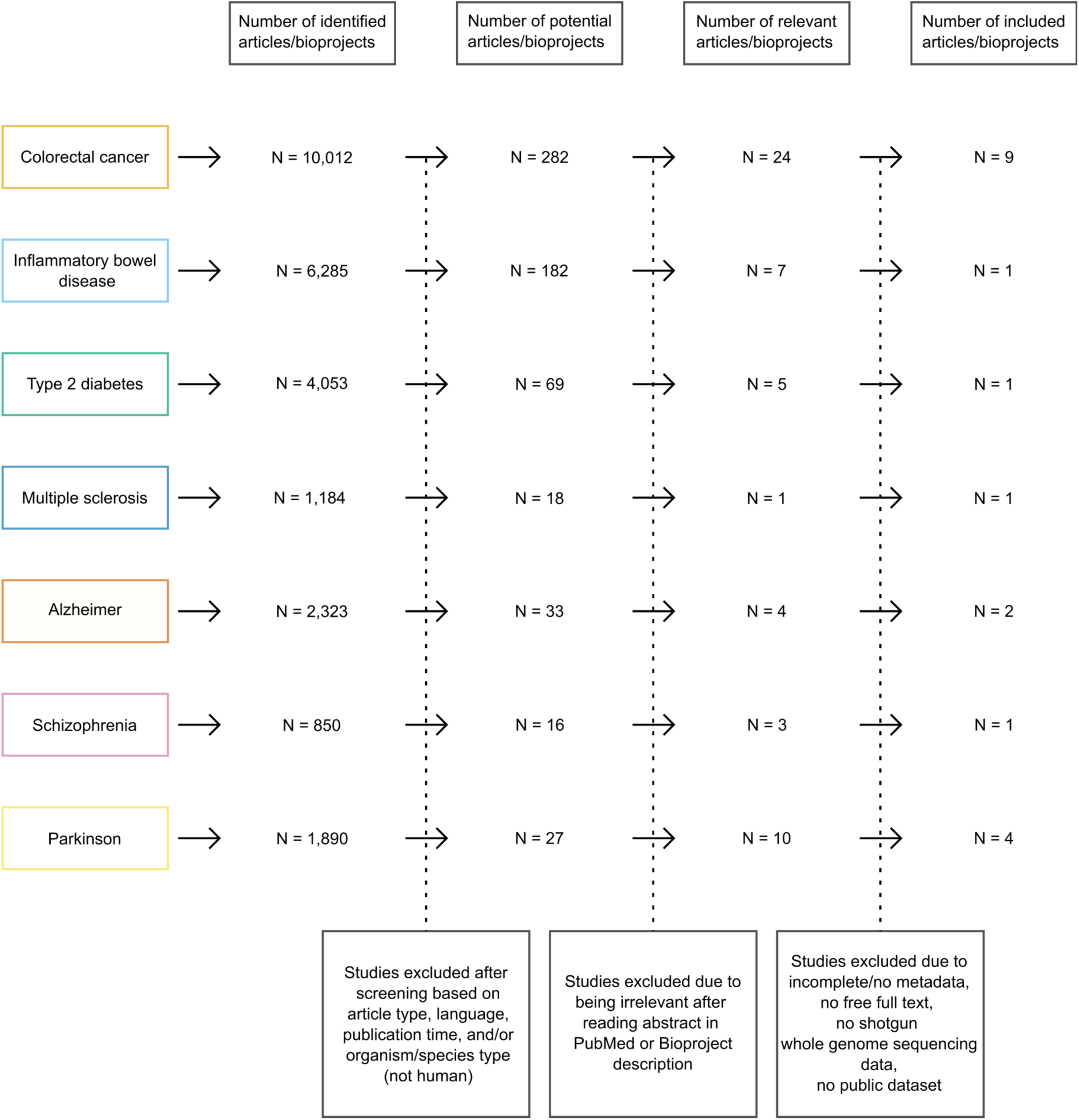
Study selection and overview of included datasets for meta-analysis. A total of 631 studies were initially screened for potential inclusion in the meta-analysis. Following eligibility assessment, 19 studies were retained, comprising 9 colorectal cancer (CRC), 1 inflammatory bowel disease (IBD), including Crohn’s disease (CD) and ulcerative colitis (UC), 1 type 2 diabetes (T2D), 1 multiple sclerosis (MS), 2 pre-Alzheimer’s disease (pre-AD; initially searched for Alzheimer’s patients, but due to the low sample sizes, we only included those with pre-AD), 1 schizophrenia (SCZ), and 4 Parkinson’s disease (PD) datasets.

### Data retrieval and processing

Raw sequencing data were retrieved from the NCBI Sequence Read Archive using SRA Toolkit (v3.1.0). Human-derived sequences were removed by aligning the raw reads to GRCh38 human reference genome using *removehuman.sh* within BBMap suite (v39.01) with default parameters. Read quality filtering, adapter trimming, base correction, and removal of low-complexity sequences were conducted concurrently using fastp (v0.23.4) with the following parameters: *-- average_qual 20, --detect_adapter_for_pe*, *--correction*, *--overrepresentation_analysis*, and *-- low_complexity_filter*. Unpaired reads resulting from trimming or filtering were corrected using BBMap (*repair.sh*), ensuring synchronization of read pairs for downstream analyses.

For microbial taxonomic profiling, we employed a Kraken2 and Bracken approach, which has been shown to outperform marker gene-based methods in recent benchmarking studies^27^. Taxonomic classification was carried out using Kraken2^28^ (v2.1.2) with the Unified Human Gastrointestinal Genome (UHGG v2.0.1) database, which comprises a total of 4,644 prokaryotic species, including 4,616 bacterial species and 28 archaeal species, and enables identification of human gut archaeal species with high resolution^26,29,30^. To improve classification accuracy and minimize incorrect lowest common ancestor (LCA) assignments, a confidence threshold of 0.3 was applied. Species-level abundance estimates were refined using Bracken (v2.7) (https://ccb.jhu.edu/software/bracken/) with read length 150. The resulting taxonomic profiles were merged into a comprehensive microbial abundance table for downstream analyses.

Given the variability in sequencing depth, read length, preprocessing methods, sample collection, and DNA extraction protocols across publicly available datasets, each dataset was processed independently. However, all were analyzed using a standardized workflow.

### Archaeome and bacteriome community analyses

Shannon diversity indices were calculated separately for archaeal and bacterial communities. Species-level abundance tables were processed in R using the phyloseq package (v1.44.0). For each dataset, samples were rarefied to the 10th percentile of library sizes to normalize sequencing depth. Shannon diversity was estimated from rarefied data using the estimate_richness() function, and comparisons between case and control groups were performed using the Wilcoxon rank-sum test. To assess beta-diversity, principal coordinate analysis (PCoA) was performed based on Bray-Curtis distance on species-level abundance profiles after normalization of counts by total sum scaling (TSS). Sample clustering patterns were visualized in two-dimensional PCoA plots, with ellipses representing group dispersion (variability of samples within a group). To statistically evaluate differences in beta-diversity, we applied permuted multivariate analysis of variances (PERMANOVA) with 999 permutations.

### Selection of CRC bacterial markers

To identify CRC-associated taxa, we conducted a PubMed search using the keywords “CRC AND bacteria AND gut AND biomarkers” for studies published until August 30, 2024. After screening relevant publications and cross-referencing additional sources, we focused on bacterial CRC biomarkers at the species level. All identified bacterial taxa were incorporated into our metabolic modeling analyses alongside *Methanobrevibacter smithii* and CRC-associated bacterial markers.

### Metabolic modeling

Draft metabolic models were generated using gapseq (development version of 1.4.0, commit acb9647) on the genomes of all CRC biomarkers namely, *Segatella copri* (formerly *Prevotella copri*) (GenBank ID GCA_020735445.1), *Mediterraneibacter torques* (formerly *Ruminococcus torques)* (GenBank IDGCA_000210035.1), *Otoolea symbiosa* (formerly *Clostridium symbiosum* (GenBank ID GCA_008632235.1), *Escherichia coli* (GenBank ID GCA_000025745.1), *Prevotella intermedia* (GenBank ID GCA_001953955.1), *Gemella morbillorum* (GenBank ID GCF_900476045.1), *Peptostreptococcus stomatis* (GenBank ID GCA_000147675.2), *Porphyromonas asaccharolytica* (GenBank ID GCA_000212375.1), *Akkermansia muciniphila* (GenBank ID GCA_009731575.1), *Fusobacterium nucleatum* (GenBank ID GCA_003019295.1), *Parviromonas micra* (GenBank ID GCA_900637905.1), *Bacteroides fragilis* (GenBank ID GCA_000025985.1), as well as *M. smithii* (GenBank ID GCA_000016525.1). The gut medium formulation used in this study followed previously established protocols^31,32^. Subsequently, the initial metabolic models were gap-filled utilizing this specified medium. To simulate co-culture interactions, community metabolic models were constructed for each pair of organisms using PyCoMo^33^ version 0.2.7. The corresponding gut medium was applied to each model. The simulations included calculations of community growth dynamics, metabolite exchanges, and cross-feeding interactions between *M. smithii* and CRC-associated bacterial biomarkers, employing PyCoMo’s default settings.

For the three CRC biomarkers with validated relevance^34^—*F. nucleatum*, *E. coli*, and *B. fragilis*— co-cultures with *M. smithii* were also simulated on relevant culture media. For this, in addition to the gut medium, the MS+BHI medium formulation was incorporated for gap-filling procedures and subsequent PyCoMo simulations, as previously detailed in our previous study^35^ (full medium composition available in the referenced publication). Metabolite exchanges and cross-feeding patterns were computed by scaling the flux bounds from flux variability analysis based on the relative abundances of *M. smithii* and the CRC-associated bacteria, as determined by qPCR analysis performed on co-cultures. The resulting cross-feeding interaction profiles, highlighting metabolites produced by one organism and utilized by the other within each co-culture, were visualized using ScyNet^36^ integrated into the Cytoscape^37^ software (version 3.10.0).

### *E. coli* isolation and confirmation

To isolate the *E. coli* strain originally designated as *E. coli*_D based on the UHGG v2.0.1 taxonomy, and now classified as *E. coli* without the "_D" designation according to GTDB release 226 and subsequent updates (https://gtdb.ecogenomic.org/), we utilized a fecal sample from a previous study in which the co-occurrence of *M. smithii* and *E. coli*_D was confirmed through both cultivation and sequencing methods^35^. Specifically, 1 ml of the fecal sample from patient 51 (P51) of the aforementioned study was streaked onto CHROMagar™ *E. coli* medium. After incubation at 37°C for 24 h, colonies displaying a characteristic intense blue coloration were selected for downstream processing.

Genomic DNA was extracted from the selected colonies using the PureLink™ Microbiome DNA Purification Kit (Invitrogen, Thermo Fisher Scientific, USA), following the manufacturer’s protocol. To confirm the identity of the isolate as *E. coli*, two sets of primers were employed: one targeting the universal *E. coli* 16S rRNA gene (EC_F: 5′–CCAGGCAAAGAGTTTATGTTGA– 3′, EC_R: 5′–GCTATTTCCTGCCGATAAGAGA–3′)^38^, and the other targeting the *adk* housekeeping gene (adkF: 5′–ATT CTG CTT GGC GCT CCG GG–3′, adkR: 5′–CCG TCA ACT TTC GCG TAT TT–3′)^39^.

PCR amplification was performed as described previously^38^, with small modifications. Briefly, PCR amplification of the *adk* gene was performed with an initial denaturation at 95 °C for 2 min, followed by 35 cycles of 95 °C for 1 min, 54 °C for 1 min, and 72 °C for 2 min. For the 16S rRNA gene, the protocol included an initial denaturation at 95 °C for 5 min, then 35 cycles of 92 °C for 1 min, 57 °C for 1 min, and 72 °C for 30 s. Both reactions concluded with a final extension at 72 °C for 5 min. PCR products of the expected size 212 bp for the 16S rRNA gene and 583 bp for *adk*, were verified via 1.5% agarose gel electrophoresis. PCR products were subsequently purified using the Purification Kit (SolGent Co. Ltd., Daejeon, Republic of Korea), according to the manufacturer’s instructions. The identity of the isolates was confirmed by direct Sanger sequencing of the purified PCR products.

### Co-culturing assays

We selected three CRC-associated bacterial strains, *F. nucleatum* (DSM 15643), enterotoxigenic *B. fragilis* (DSM 2151), and *E. coli* strain D (updated to *E. coli* in GTDB r226), for co-cultivation with *M. smithii* DSM 2375 (=ALI), due to their CRC-relevant virulence^34^. The experimental layout is illustrated in Figure 4.

Standard archaeal medium (MS medium) was prepared under anaerobic conditions as previously described^40^. Brain heart infusion (BHI) broth was prepared under anaerobic conditions in anaerobic chamber following standard protocols and flushed with nitrogen (100%). Subsequently, 10 ml of BHI broth and 20 ml of MS medium were combined in 100 ml serum bottles under anaerobic conditions. The medium was deoxygenated with nitrogen gas, supplemented with 0.75 g/l L-cysteine under anaerobic conditions, and adjusted to pH 7.0 if necessary.

The 30 ml aliquots were sealed in 100-ml serum bottles with rubber stoppers and aluminum caps, pressurized with H₂/CO₂ (4:1), and sterilized by autoclaving at 121 °C for 20 min. Before use, 0.001 g/ml yeast extract and 0.001 g/ml sodium acetate were added under anaerobic conditions. Time point designations were standardized relative to the initial inoculation of *M. smithii* ALI: t(– 1) corresponds to the time of *M. smithii* ALI inoculation (0 h); t(0) denotes 24 h post–*M. smithii* ALI inoculation, or the point where bacterial mono-cultures in BHI + MS are initiated; t(0+) denotes the point bacterial strains are introduced to 24 h post–*M. smithii* ALI inoculation (initiation of co-cultures); t(1) represents 48 h post–*M. smithii* ALI inoculation (24 h post–bacterial inoculation for co-cultures); and t(2) indicates 96 h post–*M. smithii* ALI inoculation (72 h post– bacterial inoculation).

Cultures of *M. smithii* ALI (5 ml) were initiated at t(–1) in BHI + MS medium for further co-culturing with a bacterium, to avoid the overgrowth of bacteria due to higher incubation period for archaea and lower log phase time, with a parallel *M. smithii* ALI mono-culture prepared under identical conditions. The optical densities (OD) of *M. smithii* ALI master cultures in each experiment set used to inoculate in the mono- and co-cultures at t(−1) were: 0.07 ± 0.007 for *F. nucleatum* co-cultures, 0.05 ± 0.008 for *B. fragilis* co-cultures, and 0.47 ± 0.09 for *E. coli* co-cultures. At t(0), 0.2 ml of each bacterial strain pre-cultured overnight in BHI medium (*F. nucleatum* (OD = 0.82 ± 0.05), *B. fragilis* (OD = 0.90 ± 0.04), and *E. coli* (OD = 2.12 ± 0.02)), was separately inoculated into the *M. smithii* ALI culture to establish co-cultures, while bacterial mono-cultures were concurrently initiated in BHI + MS medium.

Optical density measurements were performed on 0.6 ml samples collected from *M. smithii* ALI mono-cultures and co-cultures at t(–1), t(0), t(1), and t(2), and from bacterial mono-cultures at t(0), t(1), and t(2). For metabolomic analyses, 1 ml samples were collected from *M. smithii* ALI mono-cultures, co-cultures, and bacterial mono-cultures at t(0), t(1), and t(2). For DNA extraction and subsequent qPCR analysis, 1 ml samples were collected from *M. smithii* ALI mono-cultures at t(2), from bacterial mono-cultures at t(1) and t(2), and from co-cultures at t(0), t(1), and t(2). At t(2), endpoint analyses were conducted on both co-cultures and *M. smithii* ALI mono-cultures to assess methane production, detect F420-positive cells, and scanning electron microscopy was also conducted at t(2) for *M. smithii* ALI mono-cultures, co-cultures, and bacterial mono-cultures.

Blank BHI and BHI + MS media were incubated in triplicate for contamination monitoring, as well as for metabolomics experiments, while all inoculated cultures were performed in five biological replicates. All cultures were maintained at 37 °C with continuous shaking at 80 rpm throughout the experiments. DNA was extracted from cultures by using the PureLink™ Microbiome DNA Purification Kit (Invitrogen, Thermo Fisher Scientific, USA) according to the manufacturer’s protocols. DNA extracts were stored at −80 °C for further analysis. Culture samples for metabolomics were also stored at −80 °C until further analysis.

### CH4 measurement and fluorescence microscopy

To evaluate whether methane (CH₄) production by *M. smithii* ALI was increased during co-cultivation, CH₄ concentrations were measured in the gas phase of the culture bottles with the CH₄ sensor BCP-CH4 (BlueSens gas sensor GmbH, Germany) following the manufacturer’s instructions. Measurements were performed for both mono-cultures and co-cultures at the end of the incubation period.

Additionally, the presence and proliferation of *M. smithii* ALI were confirmed through the detection of the characteristic autofluorescence of coenzyme F420 with maximum emission wavelength of 480^41^. Fluorescence microscopy was conducted using a Zeiss Axio Imager A1 microscope (Carl Zeiss AG, Germany) equipped with a fluorescence module, as previously described^35^.

### Scanning Electron Microscopy

The cell morphologies of the mono-cultures and co-cultures were examined using a Zeiss Sigma 500 V7P scanning electron microscope (Carl Zeiss AG, Germany). For sample preparation, 2 ml of culture (including supernatant) were immediately transferred on ice to the Core Facility Ultrastructure Analysis at the Medical University of Graz (Austria) for scanning electron microscopy (SEM), where cell pellets were prepared by centrifugation at 4000 × g for 10 min and processed as previously described^35^.

### qPCR

Quantification of the absolute copy numbers of bacterial 16S rRNA and archaeal *mcrA* genes in the samples was conducted using a quantitative real-time PCR (qPCR) method with SYBR Green dye as previously described^9^. Briefly, each qPCR reaction mixture consisted of one microliter of DNA template combined with SYBR Green Supermix (Bio-Rad). The bacterial 16S rRNA genes were amplified using the primer pair 331F (5’-TCCTACGGGAGGCAGCAGT-3’) and 797R (5’-GGACTACCAGGGTATCTAATCCTGTT-3’)^42^. The thermal cycling protocol for bacterial amplification involved an initial denaturation step at 95°C for 15 seconds, followed by 40 cycles consisting of denaturation at 94°C for 15 s, annealing at 54°C for 30 s, and extension at 73°C for 40 s. For the detection and quantification of *M. smithii* ALI, primers M1F (5’-GCAATGCAAATTGGTATGTC-3’) and M1R (5’-TCATTGCGTAGTTAGGRTAGT-3’) were used, specifically targeting the *mcrA* gene (single-copy gene). The thermal cycling conditions included an initial denaturation at 94°C for 3 s, followed by 40 cycles of denaturation at 94°C for 45 s, annealing at 56°C for 45 s, and elongation at 72°C for 30 s.

The quantification cycle (Cq) values were analyzed using the regression method available in Bio-Rad CFX Manager Software (version 3.1). The absolute gene copy numbers for bacterial 16S rRNA and methanogenic *mcrA* genes were determined based on Cq values and corresponding amplification efficiencies derived from standard curves. These standard curves were established according to previously described protocols^9^ based on standard curves from defined DNA samples of *E. coli* for bacterial 16S rRNA genes and the *mcrA* gene from the *M. smithii*^43,44^.16S rRNA gene copy numbers were normalized using species-specific values from the rrnDB database (*F. nucleatum =* 5 copies, *B. fragilis =* 6 copies, and *E. coli =* 7 copies). The detection thresholds were established using the mean Cq values obtained from non-template control reactions. All qPCR assays were performed in three technical replicates, with each assay replicated across five independent biological replicates of both mono and co-cultures.

### Metabolite quantification using Nuclear Magnetic Resonance

In order to gain initial insights into the consumption/production of amino acids, metabolic activity of *M. smithii* ALI was investigated by Nuclear Magnetic Resonance (NMR) spectroscopy using *M. smithii* ALI cultures previously described in an earlier study^40^. Measurements were performed in triplicate for *M. smithii* ALI at 72, 168, and 240 h post-inoculation in MS medium supplemented with yeast extract, following the experimental protocol outlined in the same reference.

Metabolomic profiling of mono- and co-culture samples (cell+supernatant) collected for this study at different timepoints (Figure 4) was also conducted through NMR spectroscopy. For this purpose, samples were initially treated using protein precipitation by the addition of methanol to give a methanol–water mixture at a 2:1 ratio. After centrifugation, supernatants were collected and subsequently dried by lyophilization. The lyophilized extracts were then reconstituted in a sodium phosphate-buffered solution supplemented with an internal NMR reference standard, 4.6 mM 3-trimethylsilyl propionic acid-2,2,3,3-d4 sodium salt, before being transferred into NMR sample tubes.

Spectral acquisition was performed using a Bruker Avance Neo 600 MHz spectrometer, equipped with a triple resonance inverse probe, operated at a controlled temperature of 310 K. Data were captured utilizing the Carr–Purcell–Meiboom–Gill pulse sequence over 128 scans, with acquisition and initial processing conducted via Topsin 4.5 software (Bruker GmbH, Rheinstetten, Germany). Post-acquisition data treatment, involving spectral alignment and Probabilistic Quantile Normalization (PQN), was executed using MATLAB software (version 2014b, MathWorks, Natick, MA, USA).

For the precise quantification of metabolites displaying significant enhancement in signal intensities within co-cultured samples compared to mono-culture controls, integration of targeted metabolite peaks from the aligned spectra was conducted following baseline correction utilizing trapezoidal integration methods. Subsequent normalization against the proton number, specific J-coupling characteristics, and the integral of the internal standard allowed for accurate determination of metabolite molar concentrations.

### Metabolite determination using Mass Spectrometry

The blank MS+BHI medium (n = 3 replicates) as well as supernatant of the co-culture of *M. smithii* ALI and *F. nucleatum* (n = 3 replicates) were measured using the Elute PLUS LC system (Bruker, Bremen, Germany) coupled to a timsTOF Pro 2 mass spectrometer (Bruker, Bremen, Germany) with a vacuum-insulated probe heated electrospray ionization (VIP-HESI) source in both reverse phase (RP) and hydrophilic interaction (HILIC) modes.

RP separations were conducted on an Intensity Solo 2 C18 Column (100 Å; 2.0 µm; 2.1 mm × 100 mm; #BRHSC18022100, Bruker) with 0.1% formic acid (ROTIPURAN® ≥99 %, LC-MS Grade, Carl Roth, Karlsruhe, Germany) in MilliQ water as mobile phase A and 0.1% formic acid in acetonitrile (≥99.9%, HiPerSolv CHROMANORM® for LC-MS, VWR, Darmstadt, Germany) as mobile phase B. A 5 µl injection of each sample was used. The separation was carried out at a flow rate of 0.6 ml/min with a column temperature maintained at 50°C. The following gradient was applied: 0-2 min, 5% B; 2-10 min, 5-40% B; 10-11 min, 40-98% B; 11-13 min, 98% B; 13-13.1 min, 98-5% B; 13.1-15.5 min, 5% B.

HILIC separations were performed on an ACQUITY UPLC BEH Amide column (130 Å, 1.7 µm, 2.1 mm × 150 mm; #186004802, Waters) with 10 mM ammonium formate and 0.1% formic acid in MilliQ water as mobile phase A and 10 mM ammonium formate and 0.1% formic acid in acetonitrile as mobile phase B. Injection volume was set to 5 µl, the flow rate of 0.5 ml/min and the column temperature at 40°C. The gradient was as follows: 0-1 min, 100% B; 1-6 min, 100-90% B; 6-10 min, 90-75% B; 10-11 min, 75-60% B; 11-12 min, 60% B; 12-12.1 min, 50-100% B; 12.1-21 min, 100% B.

The VIP HESI source was set to default conditions: endplate offset 500 V; capillary 4500 V; nebulizer gas 2.0 bar; dry gas 8.0 l/min; dry temp 230 °C; sheath gas 4.0 l/min; sheath gas temperature 400 °C. The probe head was put at minimum distance to the front-end assembly for maximum intensity. LC-MS/MS data were acquired in both positive and negative dda-PASEF modes, for a mass range of m/z 20-1300. Default Bruker PASEF acquisition parameters for MS/MS acquisition were used: 2 ramps (12 precursors each) per cycle; resulting cycle time 0.69 s; Intensity threshold 100 counts; Target Intensity 4000 counts (signals below that threshold will be scheduled for MS/MS fragmentation more often); active exclusion activated (0.1 min; reconsider if intensity increase is at least 2-fold). Data acquisition was performed using Bruker software timsControl® and Compass HyStar® software, and data acquisition was managed using the same software. Quality control (QC) samples were run every ten injections in HILIC mode and every five injections in RP mode, and blank samples were analyzed at the beginning and the end of each batch using H₂O for RP and methanol for HILIC.

Raw data was processed using MetaboScape® (version 2024b, Bruker, RRID:SCR_026044) with four-dimensional (4D) feature extraction, capturing mass-to-charge ratio (m/z), isotopic pattern quality, retention times, MS/MS spectra, and collision cross-section (CCS) values. Feature extraction was performed using the T-ReX® 4D algorithm (RRID:SCR_026044), followed by annotation through the Bruker Human Metabolome Database (HMDB, RRID:SCR_007712) and the NIST Mass Spectral Library (RRID:SCR_014668) as well as additional “Target lists” derived from several CCS databases (Pacific Northwest National Laboratory, METLIN, MiMeDB) resulting in Level 2 annotation according to Sumner et al.^45^. Matching was performed using four parameters: m/z (accurate mass, error <5 ppm), mSigma (isotopic pattern, score <100), MS/MS match (score >600) and CCS accuracy (error <3%).

High-quality annotations for final analysis were selected through visual inspection of chromatogram and ion mobilogram peak quality and sufficient annotation scores (match for at least 2 of the 4 parameters). Duplicate annotations were resolved by removing annotations with 1) more missing values, 2) lower overall annotation quality and 3) lower CCS accuracy until only unique annotation remained.

Raw metabolite concentrations of 489 compounds (level 2 annotation) were processed with MetaboAnalyst Version 6 (www.metaboanalyst.ca). Missing values were replaced by LoDs (1/5 of the minimum positive value of each variable). Samples were normalized by PQN to account for different cell counts and dilution effects and log transformed (base 10).

### Statistical analysis

All statistical analyses were conducted using R (version 4.3.1) in RStudio (version 2023.06.1+524). In datasets where samples were not initially matched for potential confounding variables (age, sex, and BMI), additional control for these factors was applied where feasible. Specifically, samples were matched within each dataset by sex, age (±5 years), and BMI (±3 units) to reduce potential bias (Table 1).

Subsequent differential abundance analyses for both bacteria and archaea were carried out following centered log-ratio (CLR) transformation of the dataset, using Wilcoxon rank-sum test, followed by Benjamini–Hochberg false discovery rate (FDR) correction for multiple testing. Following previously established criteria^46^, provided that more than two datasets were available, a taxon was associated with a given disease if it demonstrated a statistically significant association (*p*-adjusted < 0.05) in the same direction across at least two independent studies of the disease.

Welch’s t-test and the Wilcoxon rank-sum test were used to compare co-cultures and mono-cultures across individual NMR metabolomic profiles, qPCR data, and methane measurements based on normal distribution of data. For NMR metabolomic data, multiple testing correction was applied using the Benjamini–Hochberg FDR method. Significant changes in metabolites measured with mass spectrometry in the co-culture of *M. smithii* ALI and *F. nucleatum* compared to the blank medium were determined by t-test with a threshold of *p-*adjusted < 0.05 and minimum fold change threshold of FC > 1.5.

## Results

### Systematic reprocessing of human gut metagenomes enables standardized archaeal profiling across diverse studies

We systematically collected, reprocessed, and reanalyzed raw microbiome datasets, selecting only those studies that provided publicly accessible metagenome sequencing data (in FASTQ or FASTA format) derived from stool samples, accompanied by disease metadata (i.e., case versus control classifications) for at least 20 subjects per category. Out of an initial pool of 627 studies, 573 were excluded after abstract screening, resulting in 54 eligible studies. Subsequent refinements led to the exclusion of an additional 35 studies due to missing metadata, inconsistencies between sample identifiers in metadata and sequencing files, unavailable data or full texts, or the use of 16S rRNA amplicon sequencing instead of metagenomic shotgun sequencing, the latter being essential for our meta-analysis due to its superior species-level resolution for Archaea. Ultimately, 19 studies were retained for meta-analysis (Figure 1A, Figure S1).

The dataset initially comprised one fecal metagenome sample each for 2,881 subjects. For the study by Zhou^47^, only individuals who had not received any medication for MS treatment were included in the analysis to minimize confounding effects^26^. Additionally, for datasets where antibiotic use was explicitly reported in the metadata (e.g., Franzosa’s study^48^), we excluded those samples to avoid potential microbiome alterations due to antibiotic exposure. To minimize confounding effects of sex, age, and BMI, factors known to influence both the archaeome and microbiome^49–52^, we matched samples separately within each study whenever possible. In the studies conducted by Yu and Jo^53,54^, sample matching based on these covariates was not feasible due to incomplete metadata. However, both studies reported no significant differences in sex, age, or BMI between case and control groups. In contrast, datasets from Feng, Boktor, and Bedarf^55–57^ had already been pre-matched for sex, age, and BMI. In Franzosa’s study^48^, matching was performed solely by age, as BMI and sex information were not available. For all remaining 13 studies, we applied matching based on sex, age (±5 years), and BMI (±3 units), resulting in a final dataset of 1,882 samples (Table 1, Figure 1B).

A detailed summary of the included studies, covering study design, geographic origin, sample sizes, processing methods, and sequencing protocols, is provided in Table 1. While the studies utilized different DNA extraction kits, sequencing platforms, and sample accession numbers, each study maintained methodological consistency within its own framework. To ensure consistency, metagenomic data from all included studies were uniformly processed following a standardized protocol, and each study underwent independent analysis to avoid the batch effect (Figure 1C).

### Alpha and beta diversity of the gut archaeome is cohort- and disease-dependent

To investigate differences in archaeal composition between disease (case) and control groups, we performed a comprehensive re-analysis of multiple independent datasets. This approach enabled the identification of both shared and disease-specific archaeal taxa across different conditions.

To investigate the relationship between gut metagenome archaeal composition and disease, we assessed the beta diversity of the samples using Bray-Curtis distance to quantify microbial community dissimilarities, and principal coordinate analysis (PCoA) was employed to visualize clustering patterns based on species abundances derived from metagenomic shotgun sequencing. Associations between archaeome beta diversity and disease states were identified in CRC and pre-AD. Among CRC studies, 4 out of 9 (conducted by Feng^57^ (R^2^ = 0.029, *p* = 1E-03), Yu^54^ (R^2^ = 0.036, *p* = 4E-03), Thomas^63^ (R^2^ = 0.064, *p* = 1.4E-02), and Gupta^64^ (R^2^ = 0.368, *p* = 9E-03)) showed significant divergence. For pre-AD, both studies analyzed (Laske^66^, R^2^ = 0.037, *p* = 4E-02; Ferreiro^67^, R^2^ = 0.040, *p* = 1E-02)) demonstrated archaeal compositional differences (Figure S2). No detectable archaeal community variations were found when comparing cases to controls for CD, UC, T2D, SCZ, or PD (Figure S2).

Interestingly, in CRC studies where archaeal beta diversity exhibited significant case-control differences, bacterial beta diversity followed a similar trend (Figure S2 and Figure S3), suggesting a potential relationship between archaeal and bacterial communities in CRC within the gut, while this trend was not observed in pre-AD. Moreover, in other datasets, including two CRC studies (Zeller^59^ and Wirbel^62^), IBD (both UC and CD), three PD studies (Wallen^69^, Jo^53^, Boktor^55^), and SCZ study, bacterial beta diversity showed distinct clustering between cases and controls, whereas archaeal beta diversity remained unchanged.

Overall, these findings suggest that archaeal beta diversity differences in disease states are not consistently observed across studies. This variability may stem from the lower relative abundance of archaea, which limits statistical power, or from the possibility that archaea are less responsive to disease-related microbiome shifts compared to bacteria in these datasets. Furthermore, differences in cohort composition, geographic and environmental factors, patient individuality, dietary habits, different DNA extraction protocols as well as variances in sequencing methodologies may have contributed to the inconsistencies observed across studies, potentially influencing archaeal beta diversity outcomes^24^.

**Figure S2.**
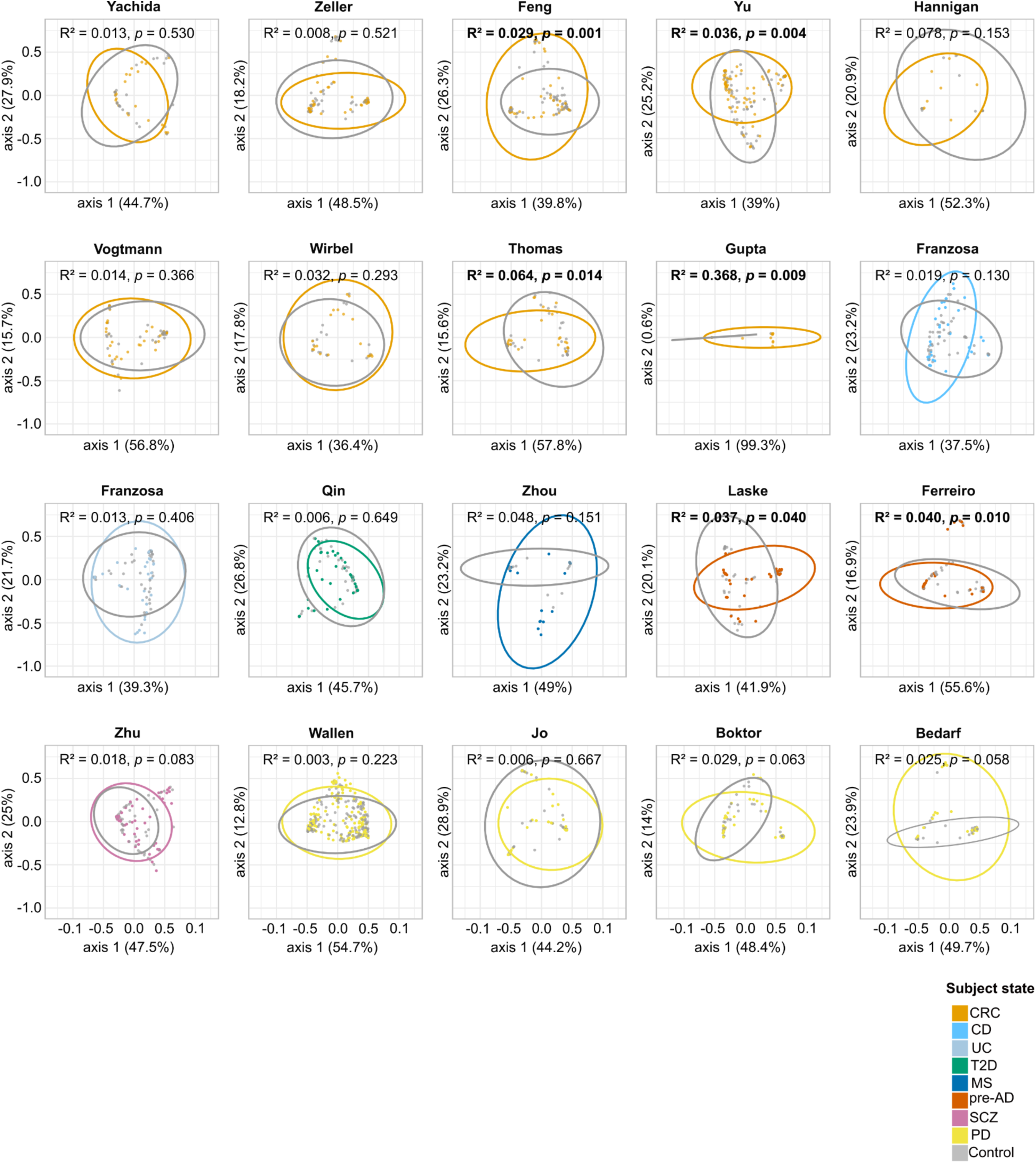
Variation in archaeal community structure across study populations, assessed by Principal Coordinate Analysis (PCoA) using Bray-Curtis distance to compare control and diseased subjects. Beta-diversity was assessed using species-level abundance profiles normalized by total sum scaling (TSS). Group variability was visualized with ellipses, and statistical differences were tested using PERMANOVA (999 permutations). Significant *p*-values (*p*-values < 0.05) are shown in bold. CRC, Colorectal cancer; CD, Crohn’s disease; UC, Ulcerative Colitis; T2D, Type 2 Diabetes; MS, Multiple Sclerosis; pre-AD, pre-Alzheimer’s Disease; SCZ, Schizophrenia; PD, Parkinson’s Disease.

**Figure S3.**
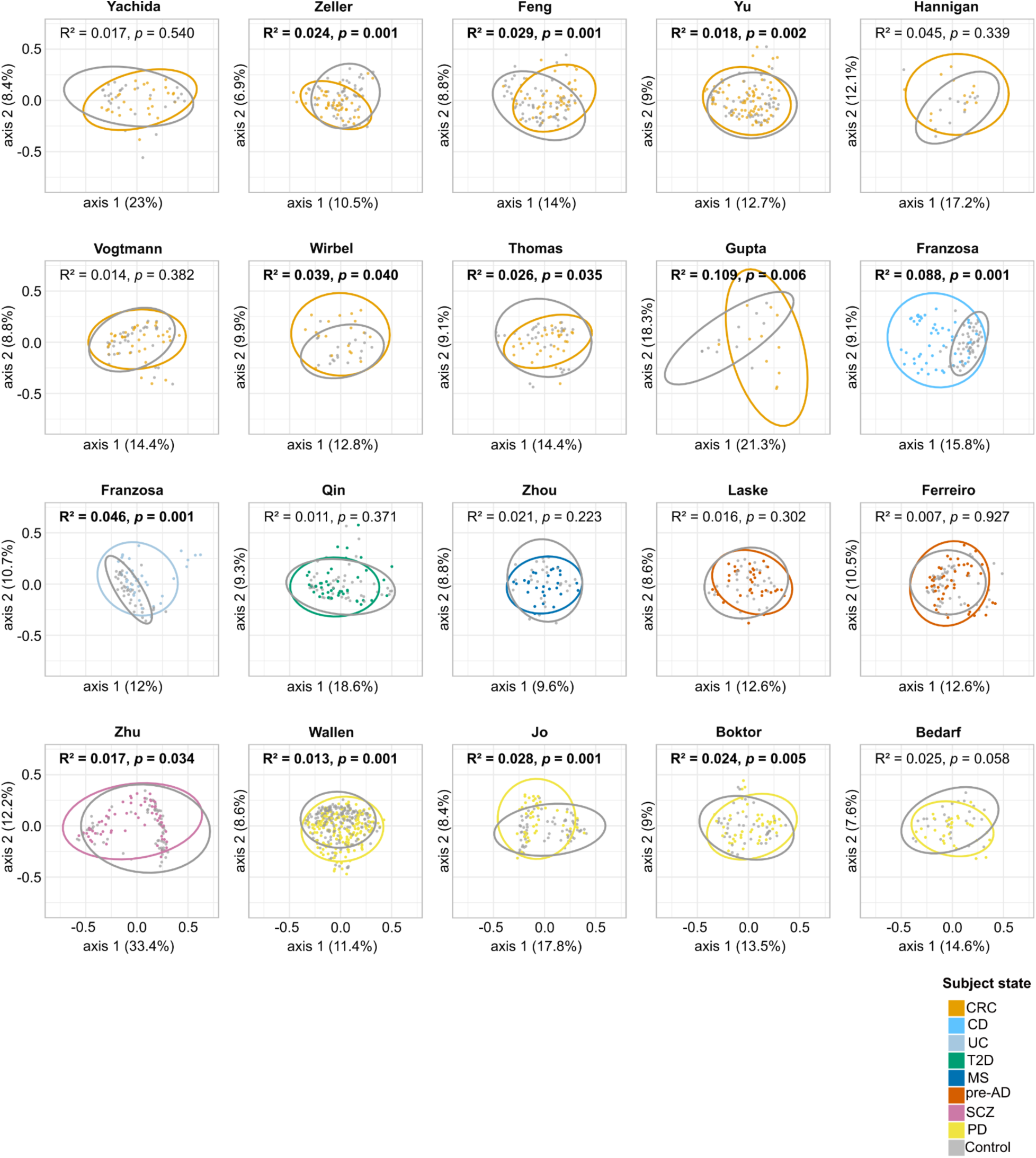
Variation in bacterial community structure across study populations, assessed by Principal Coordinate Analysis (PCoA) using Bray-Curtis distance to compare control and diseased subjects. Beta-diversity was assessed using species-level abundance profiles normalized by total sum scaling (TSS). Group variability was visualized with ellipses, and statistical differences were tested using PERMANOVA (999 permutations). Significant *p*-values are shown in bold. CRC, Colorectal cancer; CD, Crohn’s disease; UC, Ulcerative Colitis; T2D, Type 2 Diabetes; MS, Multiple Sclerosis; pre-AD, pre-Alzheimer’s Disease; SCZ, Schizophrenia; PD, Parkinson’s Disease.

**Figure S4.**
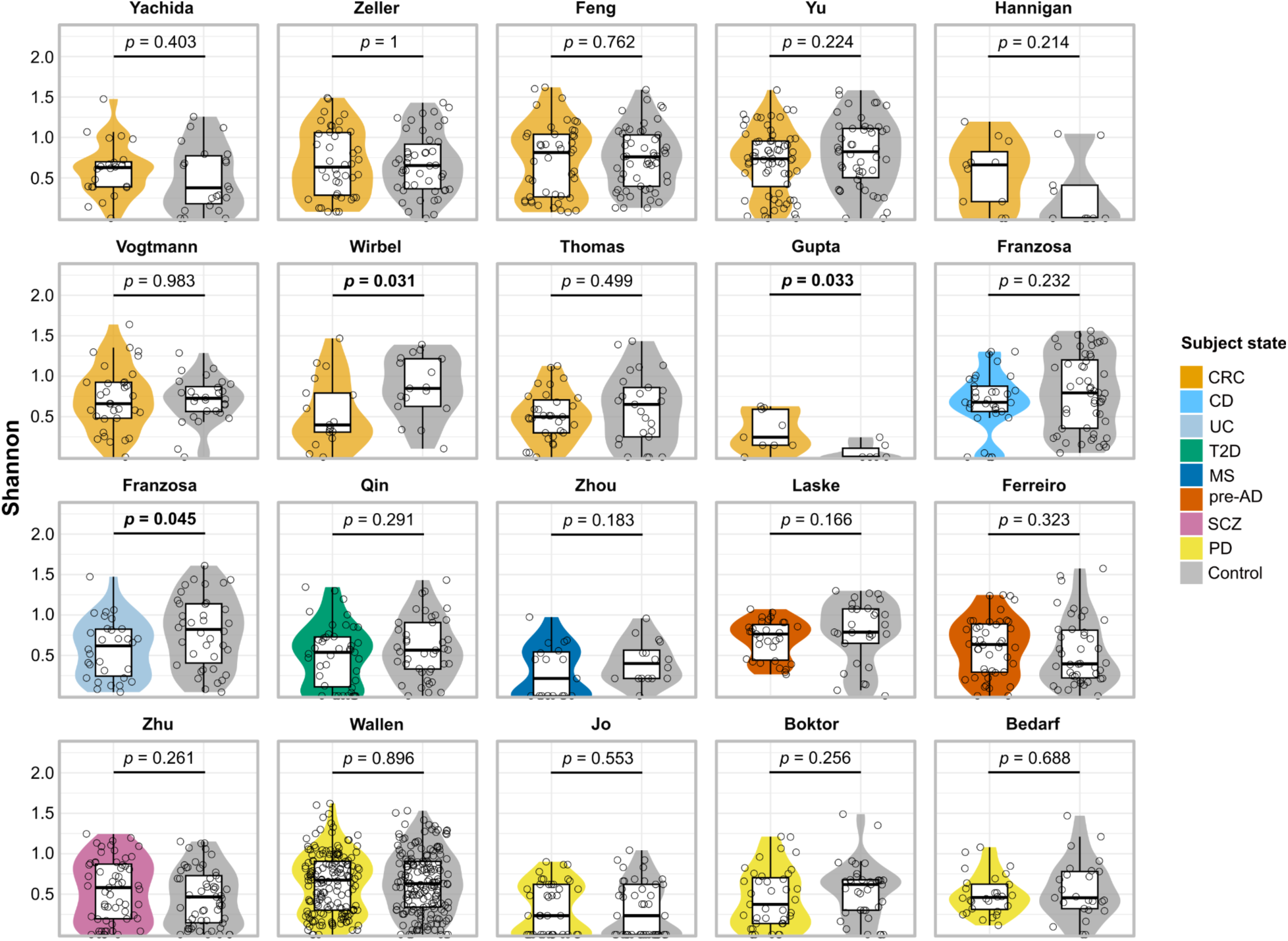
Comparison of gut archaeal diversity between disease and control groups across datasets, based on the Shannon diversity index. Case–control differences were tested with the Wilcoxon rank-sum test. Significant *p*-values (*p*-values < 0.05) are shown in bold. CRC, Colorectal cancer; CD, Crohn’s disease; UC, Ulcerative Colitis; T2D, Type 2 Diabetes; MS, Multiple Sclerosis; pre-AD, pre-Alzheimer’s Disease; SCZ, Schizophrenia; PD, Parkinson’s Disease.

Analysis of alpha diversity using the Shannon index revealed significant reductions in archaeal diversity in only one of the nine CRC studies, namely the Wirbel’s study^62^, where cases had lower diversity compared to controls (*p* = 3.1E-2), while in another study (Gupta^64^), CRC patients showed higher shannon diversity compared to the control (*p* = 3.3E-2). In contrast, UC cases (Franzosa study^48^) exhibited significantly reduced archaeal diversity compared to controls (*p* = 4.5E-2), suggesting a potential link UC and alterations in the archaeal community (Figure S4).

Notably, in studies where significant archaeal Shannon index reductions were observed, bacterial Shannon index differences followed a parallel trend, with cases showing reduced diversity compared to controls (Figure S4 and Figure S5). However, this consistent relationship was not observed across all studies of the same disease, nor in diseases for which only a single dataset was available. These findings again indicate the study-dependent shifts in archaeal diversity.

**Figure S5.**
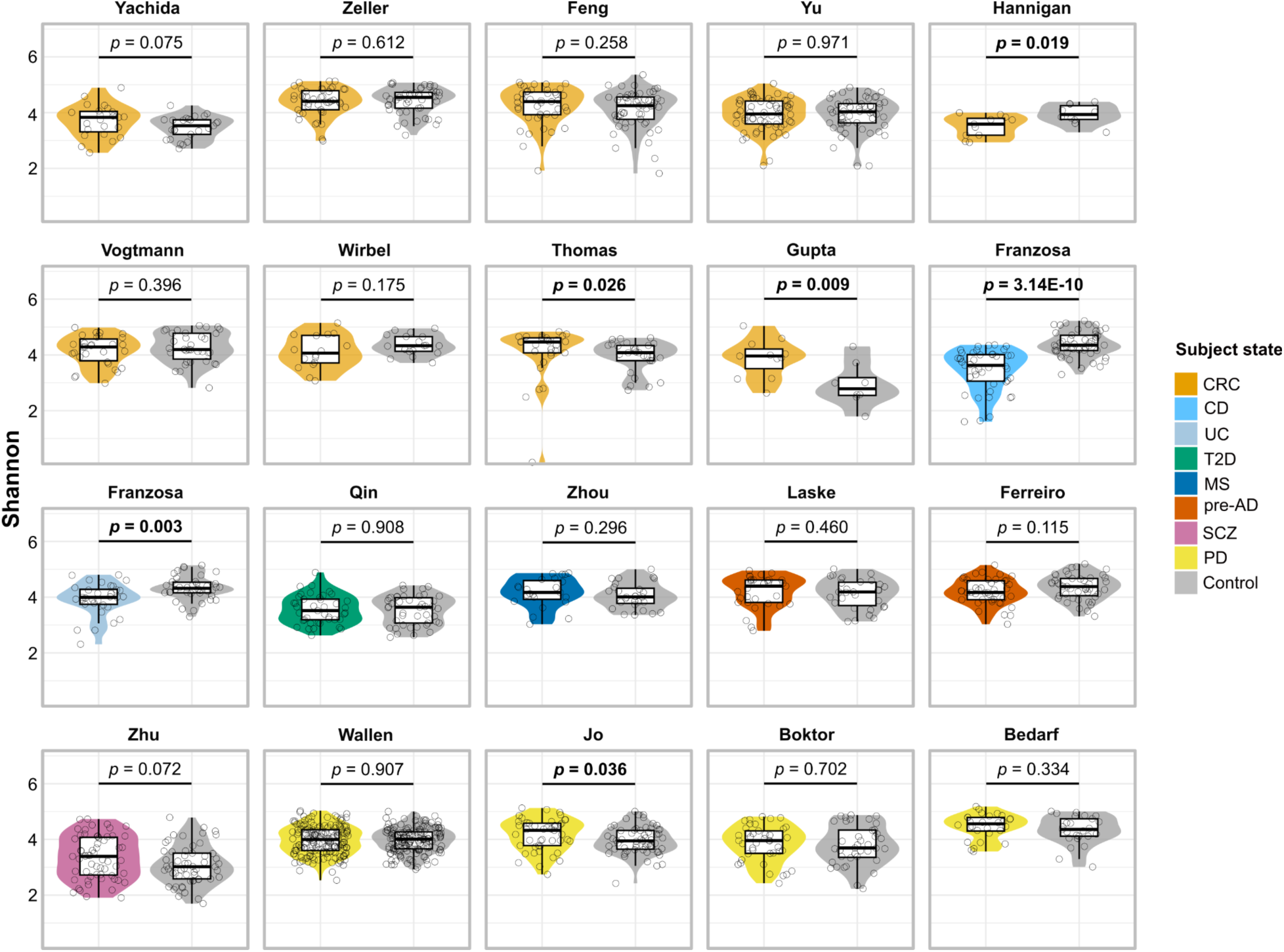
Comparison of gut bacterial diversity between disease and control groups across datasets, based on the Shannon diversity index. Case–control differences were tested with the Wilcoxon rank-sum test. Significant *p*-values (*p*-values < 0.05) are shown in bold. CRC, Colorectal cancer; CD, Crohn’s disease; UC, Ulcerative Colitis; T2D, Type 2 Diabetes; MS, Multiple Sclerosis; pre-AD, pre-Alzheimer’s Disease; SCZ, Schizophrenia; PD, Parkinson’s Disease.

### Predominant gut-associated archaea show an increasing trend in certain diseases, such as CRC

Next, we investigated if specific archaeal species are more prevalent in disease compared to the control. For each study separately (to account for study-specific confounders), we analyzed the composition of the archaeome, and compared the presence and relative abundance of archaeal species in healthy and diseased individuals.

In total, our analysis identified eight validly described archaeal species, five taxa assigned to provisional species within known genera (e.g., Methanobrevibacter_A_sp900769095), and additional genomes with low taxonomic resolution (here referred to as unclassified), defined by the Unified Human Gastrointestinal Genome (UHGG) reference catalog.

The gut archaeome was consistently dominated by various species of the genus *Methanobrevibacter*, particularly Methanobrevibacter_A_smithii (UHGG representative of *M. smithii*), Methanobrevibacter_A_smithii_A (UHGG representative of *M. intestini*), and Methanobrevibacter_A_sp900766745. These species were universally detected across all studies, regardless of the participants’ disease status. Additional *Methanobrevibacter* species, including Methanobrevibacter_A_woesei, Methanobrevibacter_A_sp900769095, and Methanobrevibacter_A_oralis, were observed at lower abundances but were consistently detected (Figure 2A). These observations suggest that members of the genus *Methanobrevibacter* are stable and persistent colonizers of the human gut, irrespective of host disease conditions.

**Figure 2.**
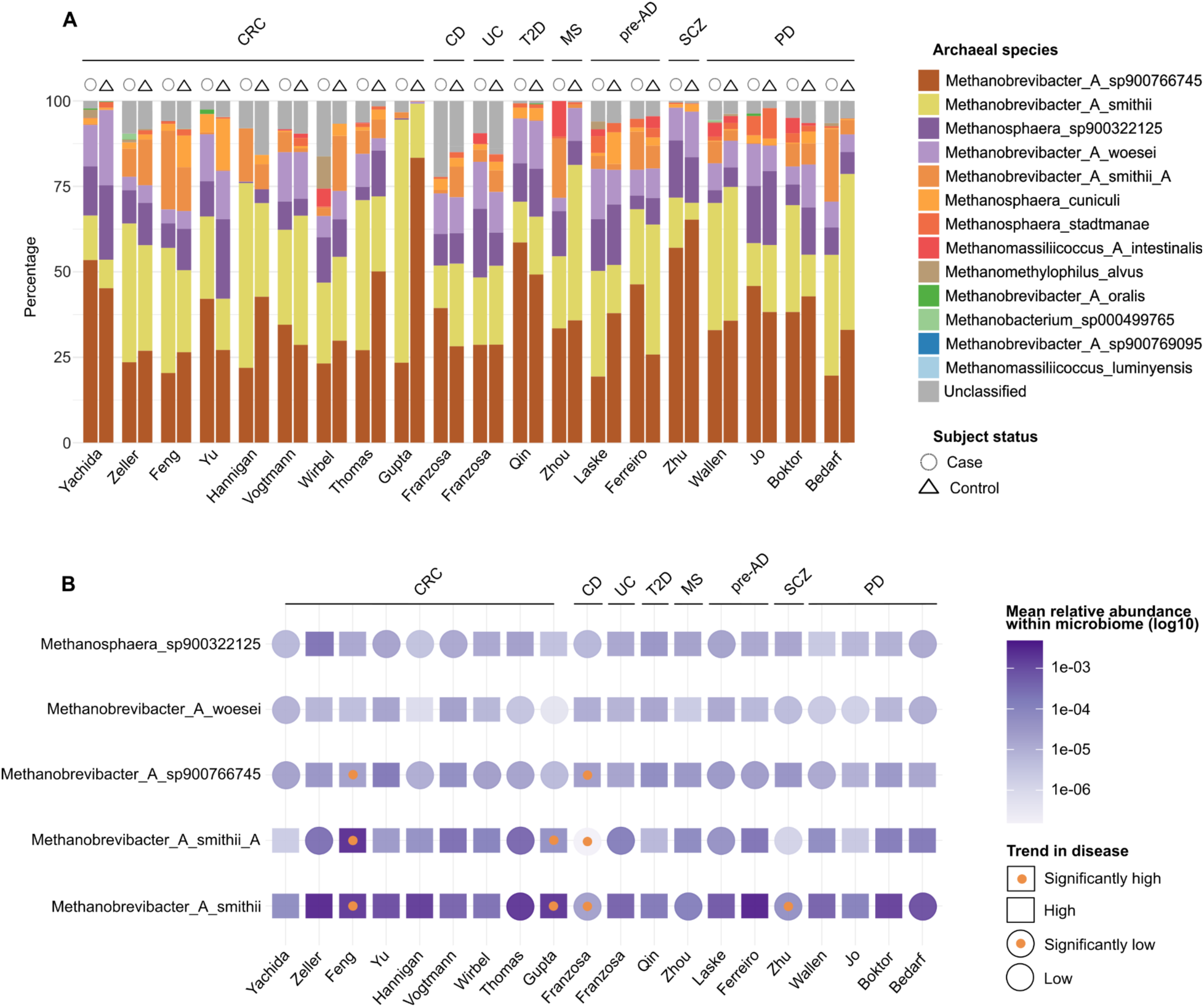
Archaeal community profiles and differential abundance analysis across disease cohorts. A) Stacked bar plots showing the relative abundances of archaeal species across shotgun metagenomic datasets, stratified by disease and study. Within each study, samples are grouped according to disease status (cases vs. controls). B) Five abundant archaeal species are represented each with a symbol indicating the direction of abundance change between disease and control groups: squares denote higher abundance in disease, and circles denote lower abundance in disease. An orange dot inside the symbol indicates a statistically significant difference (*p*-adjusted < 0.05), while empty symbols represent non-significant trends. Archaeal species are represented by their corresponding UHGG identifiers. Differential abundance was assessed using CLR-transformed data and Wilcoxon rank-sum test with FDR correction. CRC, Colorectal cancer; CD, Crohn’s disease; UC, Ulcerative Colitis; T2D, Type 2 Diabetes; MS, Multiple Sclerosis; pre-AD, pre-Alzheimer’s Disease; SCZ, Schizophrenia; PD, Parkinson’s Disease.

**Figure S6.**
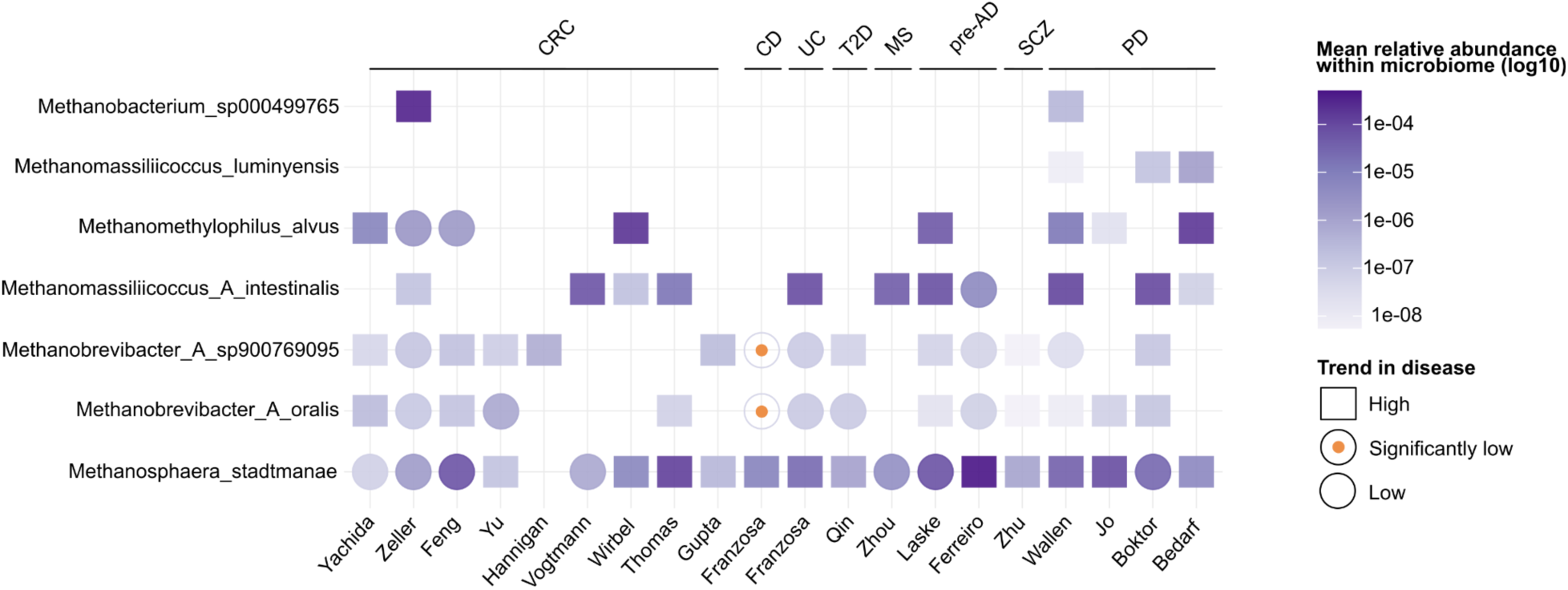
Differential abundance patterns of less dominant methanogenic archaea. differential abundance testing was performed for methanogenic archaeal species not ranked among the top five most abundant within each study. species are represented each with a symbol indicating the direction of abundance change between disease and control groups: squares denote higher abundance in disease, and circles denote lower abundance in disease. An orange dot inside the symbol indicates a statistically significant difference (*p*-adjusted < 0.05), while empty symbols represent non-significant trends. Differential abundance was assessed using CLR-transformed data and Wilcoxon rank-sum test with FDR correction. CRC, Colorectal cancer; CD, Crohn’s disease; UC, Ulcerative Colitis; T2D, Type 2 Diabetes; MS, Multiple Sclerosis; pre-AD, pre-Alzheimer’s Disease; SCZ, Schizophrenia; PD, Parkinson’s Disease.

Aside from *Methanobrevibacter*, substantial contributions to the gut archaeome included species from the genus *Methanosphaera*. Methanosphaera_sp900322125 was detected in nearly all studies and patient groups (Figure 2A).

Further diversity within the gut archaeome was represented by species from the genera *Methanomassilicoccus*, *Methanomethylophilus*, and *Methanobacterium*. Notably, Methanomassilicoccus_luminyensis was exclusively identified in PD patients, appearing in three out of four PD-related studies (Wallen^55,69^, Boktor, and Bedarf’s^56^ studies) (Figure 2A).

To explore disease-specific variations and reduce study-related bias, cases and controls were compared within individual studies. Differential abundance analysis (CLR+Wilcoxon, BH-FDR) of archaeal species were conducted in the context of the whole microbiome to account for their compositional relationships with the broader bacterial community. Among the five most prevalent archaeal species, Methanobrevibacter_A_smithii demonstrated significant (CLR+Wilcoxon, *p*-adjusted < 0.05) differences across multiple studies (Tables S1 and S2).

A potential association of *Methanobrevibacter* was observed for neurological disorders. Methanobrevibacter_A_smithii exhibited significant enrichment in SCZ patients (*p-*adjusted = 2.10E-02). Indeed, the association of higher abundances of *Methanobrevibacter* with cognitive impairment and schizophrenia has been reported in patients in one study^72^, whereas in healthy individuals, high methanogen phenotypes showed improved cognitive function in another study^3^.

An opposite trend was observed in CD, where several archaeal species were significantly reduced under disease conditions. This depletion included Methanobrevibacter_A_smithii (*p-*adjusted = 2.90E-03), Methanobrevibacter_A_smithii_A (*M. intestini*) (*p-*adjusted = 1.10E-03), Methanobrevibacter_A_oralis (*p-*adjusted = 2.47E-02) and Methanobrevibacter_A_sp900769095 (*p-*adjusted *=* 2.47E-02) while Methanobrevibacter_A_sp900766745 (*p-*adjusted = 1.72E-02) was shown to be higher compared to healthy controls (Figure 2B, Figure S6).

Methanobrevibacter_A_smithii was significantly enriched in CRC patients compared to controls in two studies (Feng^57^, *p-*adjusted *=* 1.5E-03; Gupta^64^, *p-*adjusted = 4.85E-03) (Figure 2B). In general, except for Thomas’s study^63^, Methanobrevibacter_A_smithii abundances were increased in CRC patients compared to the control group (Figure 2B).

Methanobrevibacter_A_smithii_A (*M. intestini*) similarly displayed significant enrichment in CRC patient groups in studies by Feng^57^ (*p-*adjusted = 2.61E-03) and Gupta^64^ (*p-*adjusted = 4.93E-03). Methanobrevibacter_A_sp900766745 was also enriched in CRC patients in Feng’s study^57^ (*p-* adjusted = 3.89E-02) (Figure 2B). Considering that not all studies might have used an archaea-suitable methodology^24^, these associations are a strong indicator for a potential association of *Methanobrevibacter* species with CRC. Similar positive associations of *Methanobrevibacter* signatures with CRC have been previously reported in 16S rRNA gene-based studies^73^, and a recent meta-analysis of shotgun metagenomic data^22^.

### Metabolic cross-feeding between *M. smithii* and CRC-associated bacteria suggests functional integration

A consistent, significant, and positive association between elevated *M. smithii* abundance and CRC was demonstrated across three independent studies (studies by Feng ^57^, Yu^54^, and Gupta ^64^), and was also confirmed by a recent CRC meta-analysis^74^ and our study (Figure 2B).

Given that methanogenic archaea rely on metabolic by-products of bacterial fermentation, we aimed to explore the ecological and functional interactions between *M. smithii* and bacterial species previously associated with CRC.

We curated a set of twelve bacterial taxa identified as CRC microbial biomarkers based on a literature review (Table S3). Taxa are referred to by their original names in UHGG v.2.0.1 datasets, with updated nomenclature (GTDB r226) provided in parentheses where applicable. The selected taxa include Prevotella_copri_A (updated to Segatella_copri), Mediterraneibacter_torques, Prevotella_intermedia, Peptostreptococcus_stomatis, Porphyromonas_asaccharolytica, Parviromonas_micra, Gemella_morbillorum, Clostridium_Q_symbiosum (updated to Otoolea_symbiosa), Akkermansia_muciniphila, Escherichia_coli_D (updated to Escherichia_coli), Bacteroides_fragilis, and Fusobacterium_nucleatum.

We validated the presence and enrichment of these bacterial biomarkers in our compiled CRC dataset through differential abundance analysis. after we controlled for confounding factors, such as age, sex, and BMI. Ten bacterial CRC biomarker species exhibited significant differential abundance across at least two independent studies (CLR-transformed data with the Wilcoxon rank-sum test, *p-*adjusted < 0.05), whereas Prevotella_A_copri and Akkermansia_muciniphila each demonstrated differential abundance in only one study (Figure S7). Notably, all taxa, except Mediterraneibacter_torques, appeared at least once in our selected CRC datasets, and our differential abundance findings aligned mostly with the original articles, with some small differences, which could be due differences in sample matching (Figure S7) (Table S4).

The *M. smithii* genome encodes an extensive array of transporters (see transporter file in gapseq output of *M. smithii* in Data and Code Availability), including those for amino acids (e.g. aspartate, arginine, glutamate, glutamine, or histidine), but also organic acids (e.g. succinate), indicating that the archaeal-bacterial metabolite exchange might go well beyond H_2_ and CO_2_ transfer. Experimental cultivation of *M. smithii* ALI (mono-culture in rich MS medium, Table S5) confirmed the uptake of several amino acids (leucine, valine, isoleucine, alanine, arginine, glutamic acid, methionine, asparagine, lysine, cystine, glycine, threonine, tyrosine, histidine, phenylalanine, tryptophane) and other compounds (butyrate, propionate, lactic acid, acetic acid etc.) from its environment (Figure S8, Table S5).

To functionally explore archaeal-bacterial cross-feeding potential, we performed *in silico* co-culture metabolic modeling using PyCoMo^33^. The models included all 12 bacterial CRC biomarkers (one in each simulated co-culture), and, as a control, the non-CRC-associated *Christensenella minuta*, given its known syntrophic interaction with *M. smithii*^14^.

A key finding was the universal predicted export of succinate by all bacterial partners (n=13, 100%), and the predicted import of this compound by *M. smithii* (Figure 3). This widespread pattern suggests that succinate-mediated cross-feeding could be a conserved feature of archaeal-bacterial interactions. Succinate is a common fermentation by-product during carbohydrate fermentation primarily from fumarate respiration serving for electron disposal^75,76^. *M. smithii*, encoding succinate-specific transporters, succinate dehydrogenase (Sdh) and succinyl-CoA synthetase (Suc), is thus well equipped to utilize this metabolite, potentially for redox balancing or as an anaplerotic substrate in its incomplete reductive TCA cycle^77,78^.

**Figure 3.**
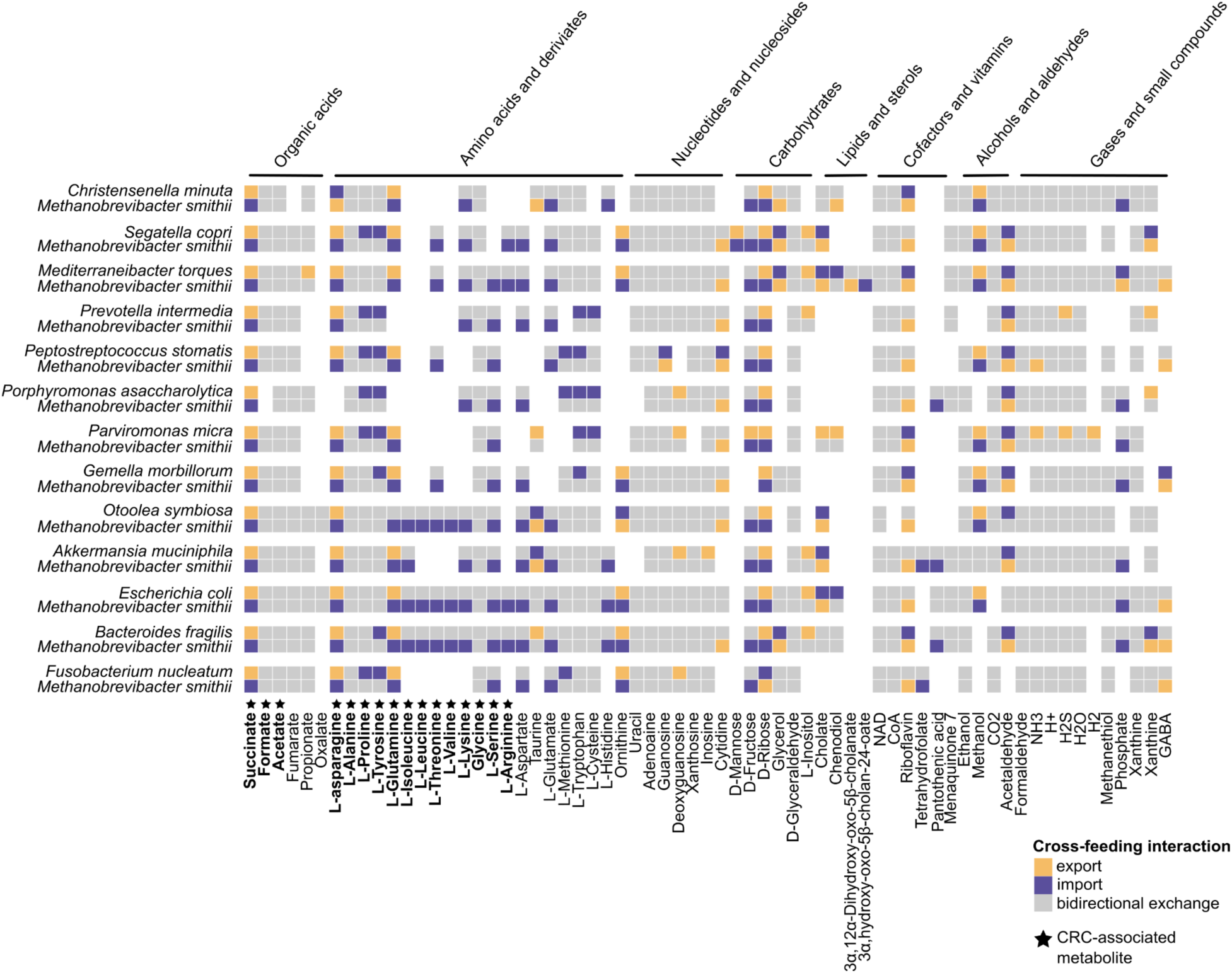
Integration of metabolic modeling analysis in colorectal cancer CRC datasets. Community-scale metabolic models show predicted cross-feeding interactions between *M. smithii* and CRC-associated bacterial biomarkers in gut medium, generated using PyCoMo. Metabolite exchanges were calculated independently of growth rates for each archaeon–bacterium pair. Colors indicate metabolite export or production (green), import or consumption (purple), or both (grey). Only cross-fed metabolites are shown. Metabolites highlighted in bold have been previously linked to CRC in the literature.

**Figure S7.**
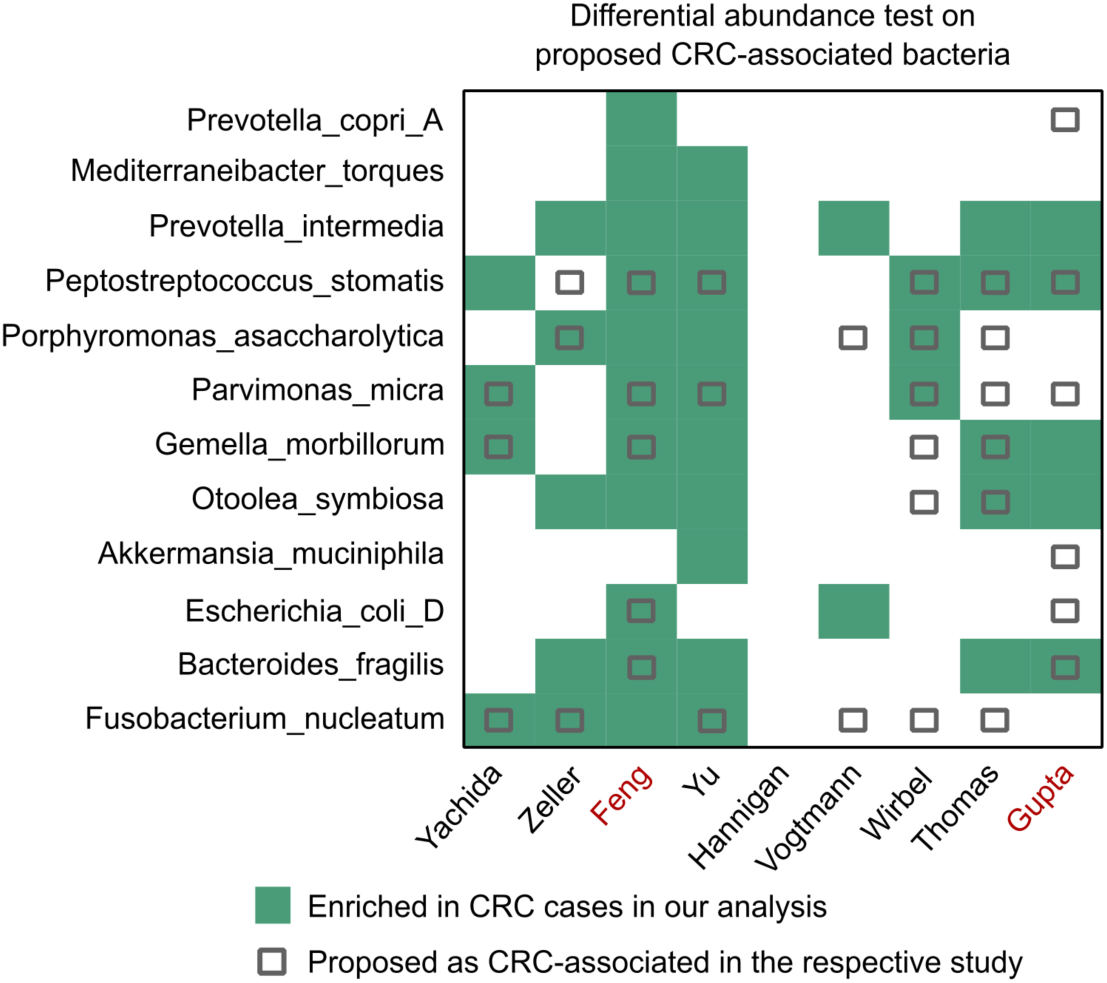
Overview of bacterial species previously reported to be associated with CRC. Green color indicates enrichment (*p*-adjusted < 0.05) in CRC based on our reanalysis of each dataset after matching for confounding variables (age, sex, and BMI). Small grey squares represent associations reported in the original publications. The study by Vogtmann et al.^61^ reported *Porphyromonas* and *Fusobacterium* as enriched at the genus level only, while Hannigan et al.^60^ focused exclusively on the virome rather than the full microbiome. Studies showing significant increase of Methanobrevibacter_A_smithii are highlighted in red. Differential abundance was assessed using CLR-transformed data and Wilcoxon rank-sum test with FDR correction.

**Figure S8.**
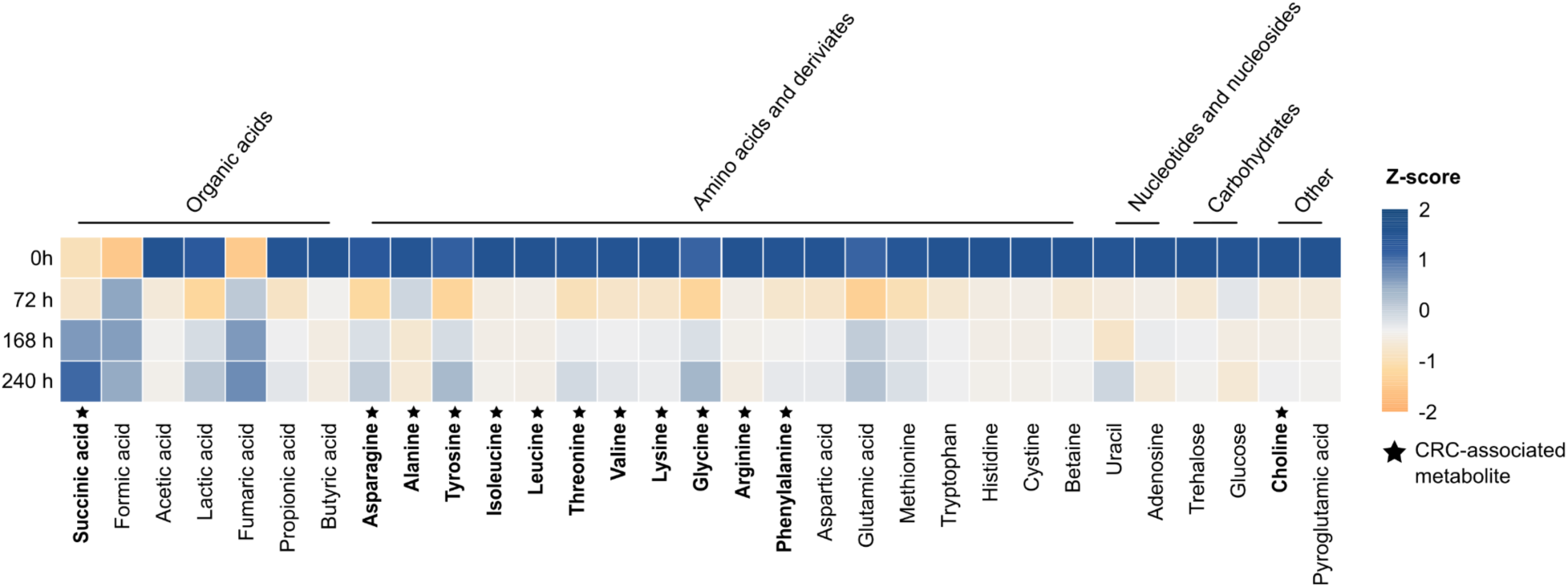
Uptake of amino acids and other compounds by *M. smithii* ALI. Metabolite analysis was performed using NMR spectroscopy on three biological replicates of *M. smithii* ALI, previously grown in MS medium and sampled at 0, 72, 168, and 240 h post-inoculation, as described in a previous study^40^. Each cell represents the Z-score of the respective metabolite, calculated across all samples as the number of standard deviations from the mean.

The relevance of succinate extends beyond microbial metabolism. Succinate, produced by bacteria such as *Fusobacterium nucleatum* has been shown to serve as a critical metabolic signaling molecule implicated in promoting CRC metastasis by activating the transcription factor STAT3, which induces epithelial-to-mesenchymal transition and enhances tumor invasiveness and metastatic potential.

Additionally, succinate suppresses the cGAS-interferon-β signaling pathway, reducing CD8+ T cell infiltration in tumors and consequently contributing to impaired anti-tumor immunity^79–81^. This positions succinate cross-feeding as a mechanistic link between microbiome metabolism and host oncogenic processes^82^.

Beyond succinate, *M. smithii* models consistently exported riboflavin in all pairwise interactions. Riboflavin is an essential precursor for the synthesis of flavin coenzymes, flavin adenine dinucleotide (FAD) and flavin mononucleotide (FMN) which play a central role in oxidation-reduction reactions redox metabolism^83^ and consequently, *M. smithii* may be beneficial to its bacterial partners. Additional predicted export-import dynamics were observed for metabolites such as amino acids L-asparagine, L-glutamine, taurine, ornithine, various carbohydrates, methanol, and acetaldehyde (Figure 3).

Notably, in interactions with CRC-associated bacteria, *M. smithii* models exhibited uptake of amino acids, including L-asparagine*, L-glutamine*, L-leucine*, L-threonine*, L-valine*, L-lysine*, L-serine*, L-arginine*, L-aspartate, L-glutamate (Figure 3). Most of these amino acids (indicated by *) have previously been linked to CRC^79,81,84–91^. These amino acids contribute to CRC progression by supporting structural protein synthesis, providing alternative energy sources, and activating oncogenic pathways such as mTORC1 (mechanistic target of rapamycin complex 1). Their elevated levels in tumor tissues reflect increased uptake and metabolic reprogramming by cancer cells and have been shown to be further shaped by gut microbial interactions. The uptake of these amino acids by *M. smithii* suggests a potential co-feeding pattern that may not only support microbial interactions, but could also influence host metabolic or signaling pathways.

### Co-culture with *M. smithii* promotes bacterial growth and metabolite production linked to CRC biomarkers

To investigate the interactions between *M. smithii* and CRC–associated bacteria, we performed a series of controlled co-cultivation experiments under strictly anoxic conditions. We selected *F. nucleatum*, enterotoxigenic *Bacteroides fragilis*, and *Escherichia coli* and their well-documented roles in CRC pathogenesis. These bacteria are known to contribute to tumor development through specific virulence factors, including colibactin, cytolethal distending toxin (CDT), cycle-inhibiting factor (CIF), and cytotoxic necrotizing factor (CNF) in *E. coli*; the adhesin FadA in *F. nucleatum*; and the enterotoxin Bft in *B. fragilis*^34^. Notably, the *E. coli* strain, previously referred to as strain D, was isolated in our previous study from a methane-producing patient, where it was found to co-occur with *M. smithii*^35^. As archaeal representative, we chose the human gut strain *M. smithii* ALI (the type strain of *M. smithii* was isolated from an anaerobic digester).

To comprehensively evaluate growth dynamics and interspecies interactions, we employed a multi-faceted analytical framework (Figure 1D, Figure 4). Growth was monitored over time via optical density (OD) measurements and quantitative PCR (targeting bacterial 16S rRNA and archaeal *mcrA* genes) in both mono- and co-cultures (n = 5 biological replicates per condition). To further assess physiological and morphological features, we performed fluorescence microscopy, scanning electron microscopy (SEM), methane quantification, and metabolomic profiling.

**Figure 4.**
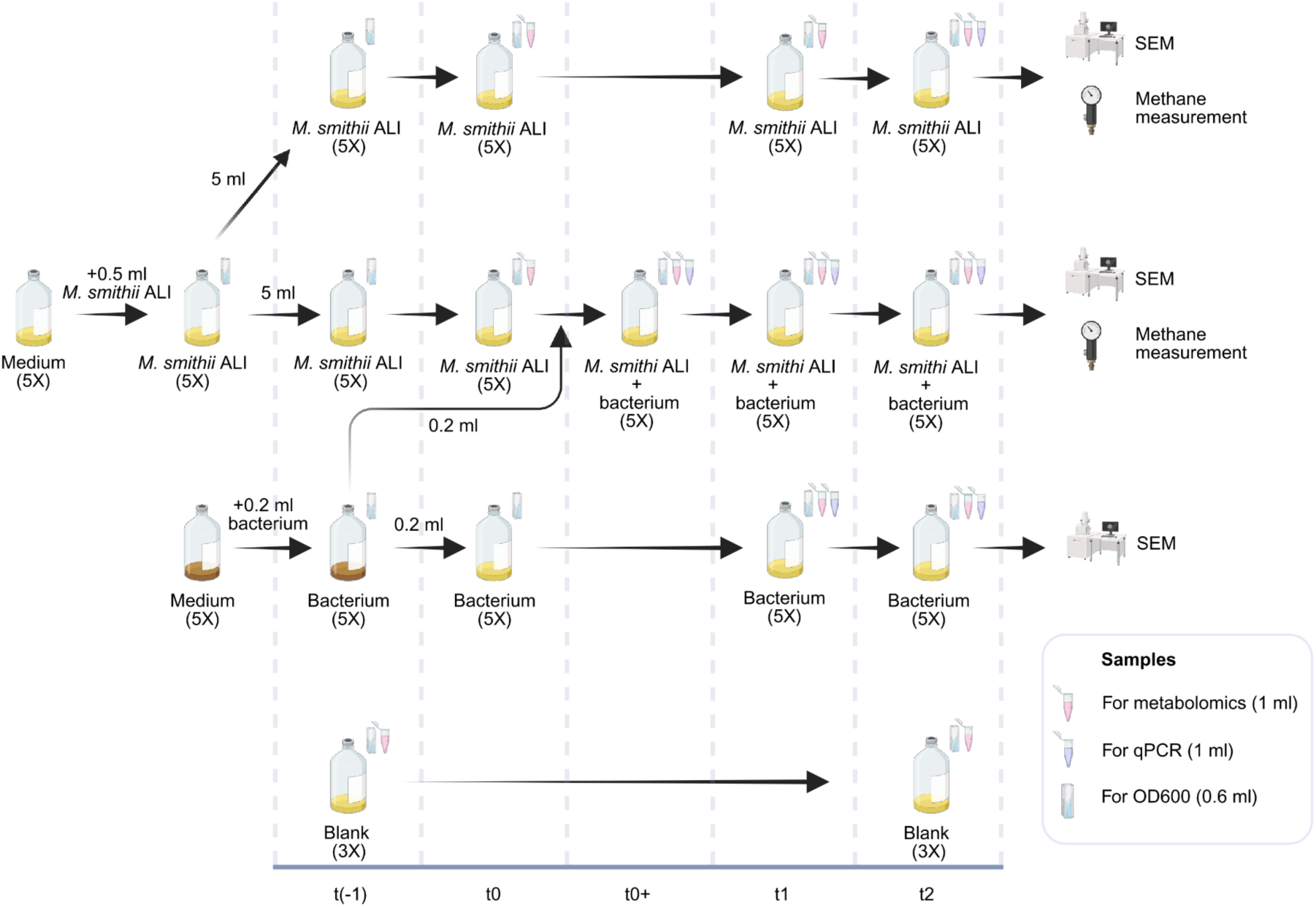
Experimental setup for co-culturing *Methanobrevibacter smithii* DSM 2375 (=ALI) with three CRC-associated bacterial strains: *Fusobacterium nucleatum* (DSM 15643), enterotoxigenic *Bacteroides fragilis* (DSM 2151), and *Escherichia coli* isolated from the fecal sample of a methane-producing subject previously shown to be co-occurring with *Methanobrevibacter smithii*^35^. Bacterial strains were first cultured overnight in BHI medium and subsequently inoculated into a combined BHI + MS medium under anaerobic conditions for co-cultivation. To account for differences in growth dynamics, 5 ml of *M. smithii* was pre-inoculated 24 h prior to bacterial addition, ensuring the archaeon had time to establish before faster-growing bacteria were introduced. Time points were defined relative to *M. smithii* ALI or bacterial inoculation: t(0) = 24 h post-inoculation, before the addition of bacteria, t(0+) = 24 h post inoculation of *M. smithii* ALI and right after the inoculation of bacterium; t(1) = 48 h post-archaeal inoculation, 24 h post-bacterial inoculation; t(2) = 96 h post-archaeal inoculation, 72 h post-bacterial inoculation. Samples were collected at key intervals for OD measurements, qPCR, metabolomics, and endpoint analyses including methane production, F420 fluorescence, and SEM. Experiments included five biological replicates for cultures and three replicates of medium control. Figure was made using BioRender.

SEM analysis revealed close proximity between *M. smithii* ALI and bacterial partners in all co-culture conditions, without showing any specific morphological alterations relative to mono-cultures. All co-cultures appeared densely populated and contained numerous archaeal and bacterial cells undergoing division (Figure 5A). Consistent with SEM observations, fluorescence microscopy revealed the presence of *M. smithii* strain ALI cells in co-cultures exhibiting the characteristic shape of this archaeal strain and F_420_-based autofluorescence in both the mono- and co-culture conditions, indicating active growth.

**Figure 5.**
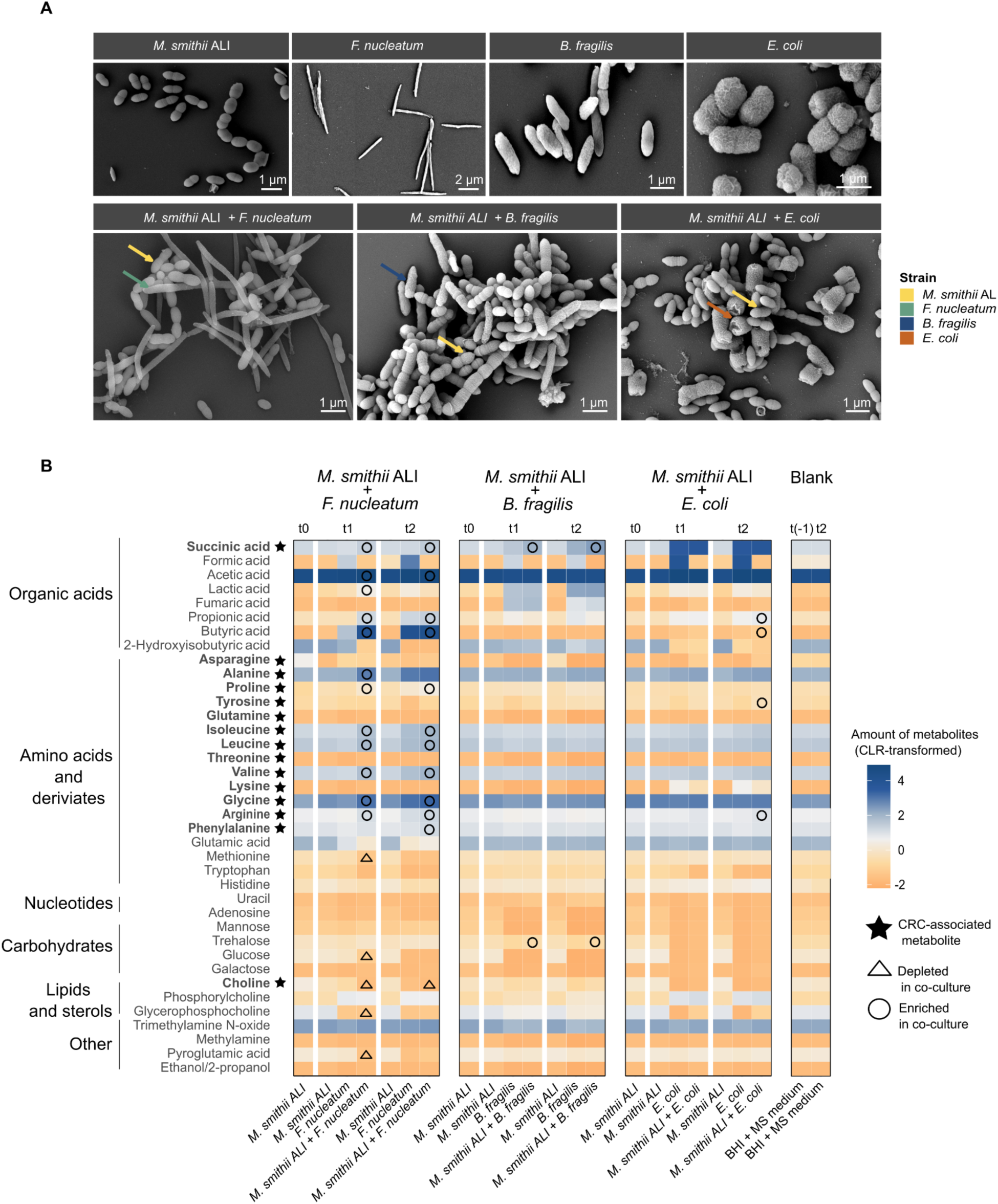
Scanning electron microscopy and NMR metabolomic profiling of *Methanobrevibacter smithii* ALI in mono- and co-culture with colorectal cancer-associated bacterial strains. A) Scanning electron microscopy (SEM) analysis of *Methanobrevibacter smithii* ALI and the bacterial strains used in co-culture experiments, including the colorectal cancer-associated species *Fusobacterium nucleatum*, *Bacteroides fragilis*, and *Escherichia coli*. Imaging was performed for both mono-cultures and co-cultures with *M. smithii* ALI. B) The heatmap displays normalized area under the curve (AUC) values obtained from metabolomics analysis, representing the relative abundance of metabolites across conditions. For improved comparability and visualization, the amount of metabolites were centered and log-ratio transformed. A metabolite was classified as enriched or depleted in co-culture only if its abundance was significantly higher or lower, respectively, compared to both *M.smithii* ALI and the corresponding bacterial mono-culture as well as in relation to the blank medium. For each co-culture experiment, mono-cultures of *M. smithii* ALI and the respective bacterial strain were grown in parallel under identical conditions. Metabolites previously associated with colorectal cancer are highlighted in bold and indicated with an asterisk. Time points are defined as follows: t(−1): initial of the experiment prior to inoculation; t0: 24 h post-inoculation of *M. smithii* ALI; t1: 24 h post-inoculation of the bacterial strain (48 h post *M. smithii* ALI inoculation) t2: 72 h post bacterial inoculation (96 h post *M. smithii* ALI inoculation). A blank medium control was included to account for background metabolite levels. Metabolite depletion (triangle) or enrichment (circle) in co-cultures was determined using Wilcoxon rank-sum test, adjusted-*p* < 0.05.

**Figure S9.**
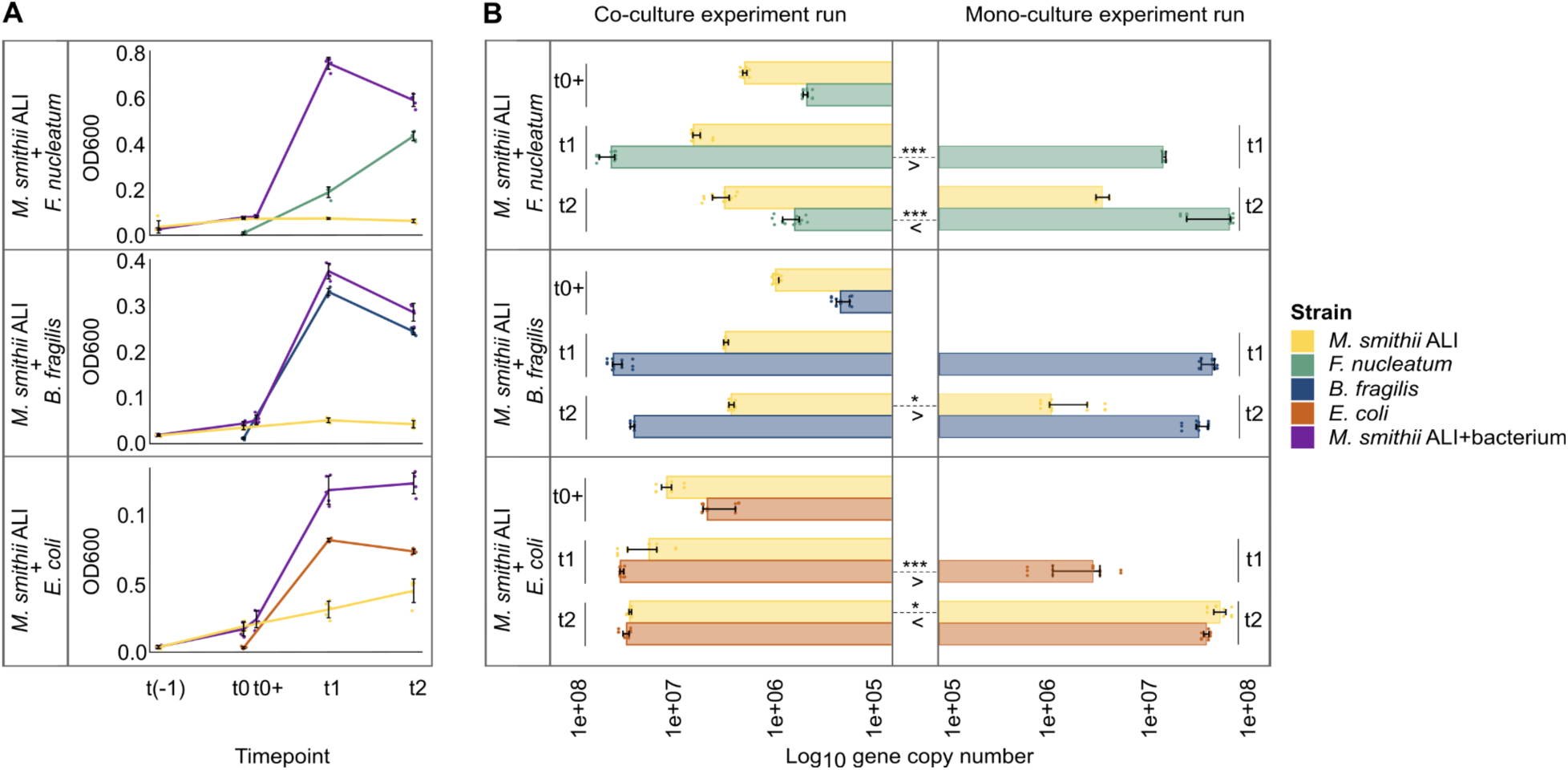
Growth dynamics and quantification of *Methanobrevibacter smithii* ALI and CRC-associated bacteria in mono- and co-culture setups. A) Optical density (OD) measurements of *M. smithii* ALI, *F. nucleatum*, *B. fragilis*, and *E. coli* grown individually and co-culture of these bacterial strains with *M. smithii* ALI. B) Quantification of *M. smithii* ALI using *mcrA* gene copy number and of CRC-associated bacterial species (*F. nucleatum*, *B. fragilis*, and *E. coli*) using 16S rRNA gene copy number in mono- and co-culture conditions. Bacterial 16S rRNA gene copy numbers were normalized using species-specific values from the rrnDB database (*F. nucleatum =* 5 copies, *B. fragilis =* 6 copies, and *E. coli =* 7 copies). For each co-culture setup, mono-cultures of *M. smithii* ALI and the respective bacterial strain were grown in parallel under identical conditions. Time points were defined as t1: 24 h post bacterial inoculation (48 h post *M. smithii* ALI inoculation), and t2: 72 h post bacterial inoculation (96 h post *M. smithii* ALI inoculation). Each dot represents a single biological replicate. \**p* < 0.05; \*\*\**p* < 0.001.

**Figure S10.**
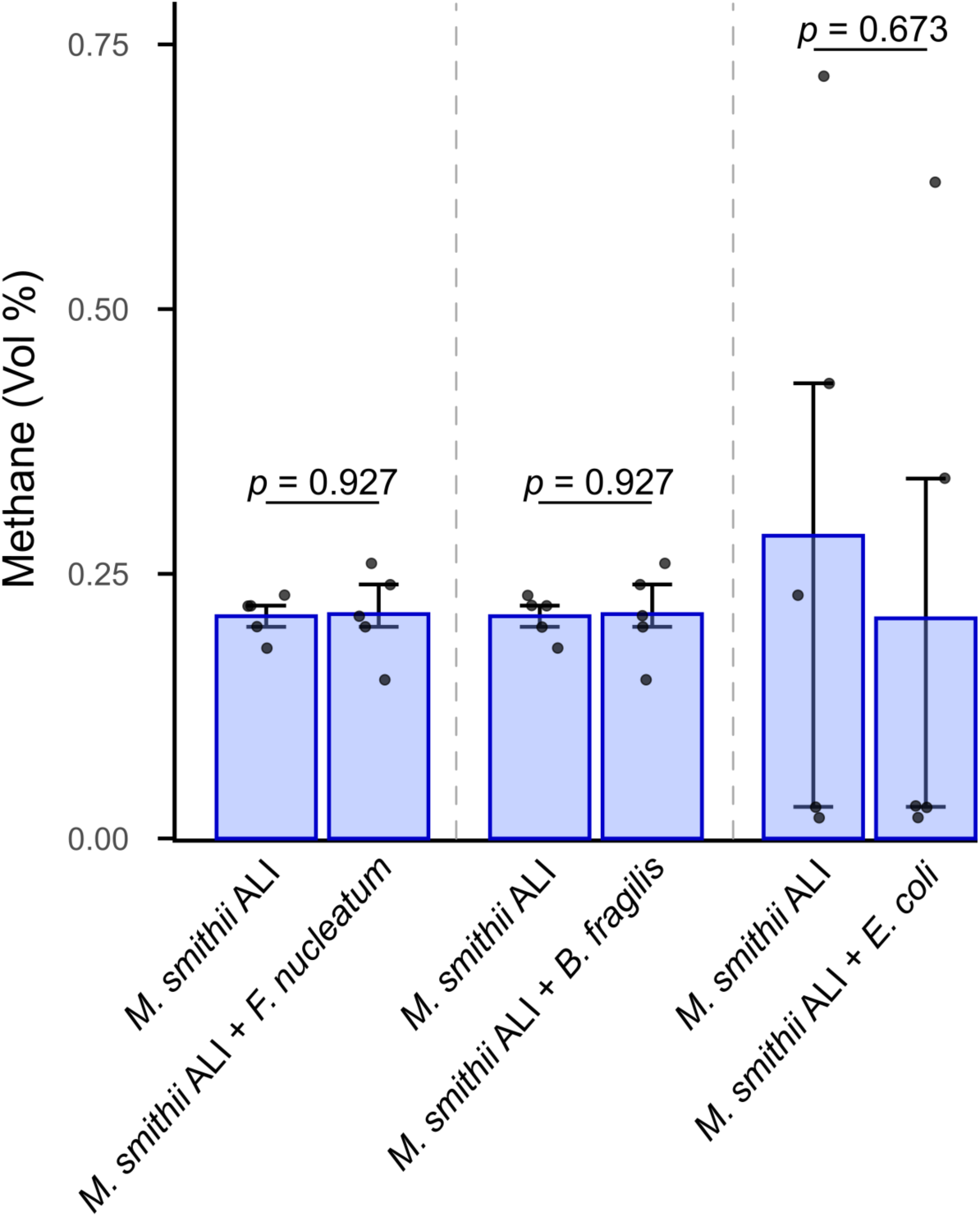
Methane concentrations produced by *Methanobrevibacter smithii* ALI are shown for both mono-culture and co-culture conditions with the respective bacterial strain. Data points represent measurements from five biological replicates, and error bars indicate standard deviations. For each co-culture experiment, mono-cultures of *M. smithii* ALI and the corresponding bacterial strain were cultivated in parallel under identical conditions to allow direct comparison of methane production dynamics.

**Figure S11.**
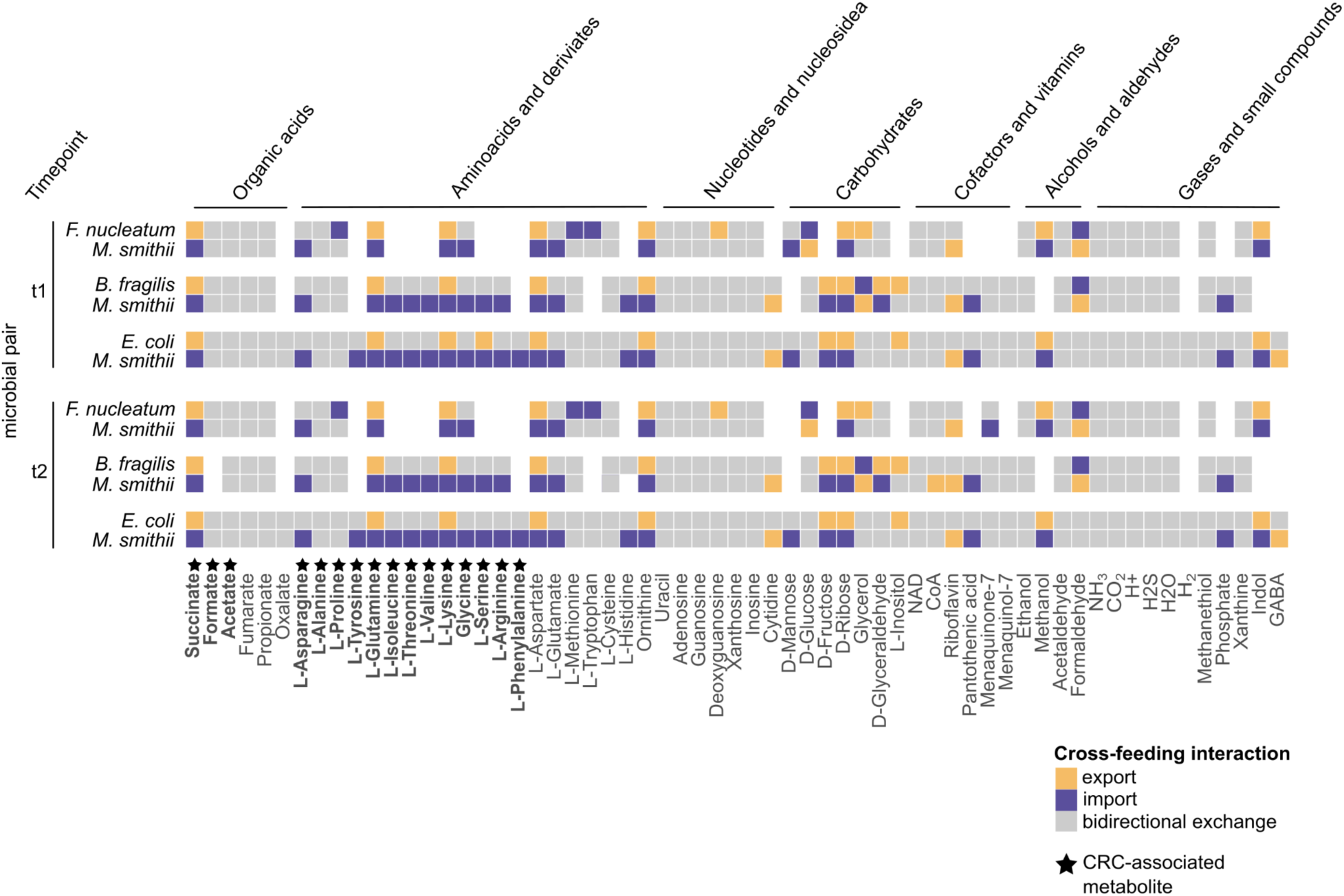
Predicted metabolite exchanges were based on relative abundances of each archaeon–bacterium pair in co-culture at two time points (T1 and T2), as determined by qPCR. At t1, mean relative abundances were: *F. nucleatum* (=0.8845) and *M. smithii* ALI (=0.1155); *B. fragilis* (=0.921) and *M. smithii* ALI (=0.0780); *E. coli* (=0.6262) and *M. smithii* ALI (=0.3738). At T2: *F. nucleatum* (=0.9274) and *M. smithii* ALI (=0.0726); *B. fragilis* (=0.9840) and *M. smithii* ALI (=0.0160); *E. coli* (=0.5696) and *M. smithii* ALI (=0.4304) (Table S6). Arrows represent predicted exchange of cross-fed metabolites, with color indicating directionality: export (green), import (purple), and bidirectional exchange (grey). Only metabolites exchanged between species are shown. Metabolites highlighted in bold have previously been implicated in CRC according to the literature (REF).

Co-culture dynamics were highly specific. *M. smithii* ALI exhibited increased growth (*p* = 4.1E-02) (evidenced by elevated *mcrA* gene copy numbers) only in the presence of *B. fragilis* compared to mono-culture conditions. *F. nucleatum* (*p* = 3.07E-06) and *E. coli* (*p* = 4.61E-06) displayed significantly enhanced 16S rRNA gene copy numbers at time point 1 compared to mono-cultures, indicating archaeon-mediated stimulation of bacterial growth speed (Figure S9). However, methane production by *M. smithii* ALI was not significantly different between mono-cultures and none of the co-cultures (Figure S10). Notably, none of the co-cultures resulted in a statistically significant mutual growth benefit, suggesting asymmetrical or unidirectional metabolic support (Figure S9).

Metabolic profiling further elucidated the biochemical landscape of the archaeal-bacterial interactions. In agreement with *in silico* modeling, all co-cultures produced higher amounts of succinate compared to their mono-cultures, confirming this metabolite as a conserved cross-feeding intermediate (Figure 3, Figure 5B, Figure S11). While metabolic alteration in *B. fragilis* and *E.coli* co-cultures with *M. smithii* ALI were relatively modest beyond succinate, the *F. nucleatum M.smithii* pairing exhibited a distinct and expansive metabolic signature.

Specifically, co-cultivation with *F. nucleatum* led to a significant accumulation of succinic acid*, acetic acid, lactic acid, propionic acid, butyric acid, and a range of amino acids, including alanine*, proline*, isoleucine*, leucine*, valine* and glycine*, arginine*, phenylalanine*. Conversely, notable consumption of methionine, glucose, choline, glycerophosphocholine, and pyroglutamic acid was observed (Tables S7, S8, and S9). Several of these amino acids (marked with *) have previously been implicated in metabolism of CRC ^79,81,84–91^, reinforcing their potential functional relevance.

Although the co-culture results did not entirely recapitulate the *in-silico* predictions (e.g. the predicted production of asparagine and glutamine could not be measured; Figure 5B and Figure S11), both approaches converged on the identification of succinate and amino acids as key mediators of archaeal-bacterial metabolic exchange. These findings indicate *M. smithii* as an active participant in CRC-associated microbial networks, potentially shaping the tumor microenvironment through metabolite cross-feeding.

### CRC-associated compounds are prominent in the *F. nucleatum*–*M. smithii*

### symbio-metabolome

To further investigate the metabolites present in co-culture, we subjected the supernatant from three replicates of *M. smithii* ALI and *F. nucleatum* co-culture at time point t1 (corresponding to the highest observed growth, as shown in Figure S9) to detailed mass spectrometry (MS) analysis. This analysis aimed to identify metabolites specifically enriched under co-culture conditions compared to the blank control medium (MS + BHI).

The co-culture supernatant displayed a diverse range of metabolite classes, including organic acids, amino acids and their derivatives, nucleotides, carbohydrates, lipids, vitamins, signaling molecules, and peptides (Figure 6, Table S10). Notably, several amino acids previously associated with CRC, such as phenylalanine and valine, as well as amino acid derivatives, including N-acetyl-tyrosine, N-acetyl-valine, N-acetyl-phenylalanine, N6-acetyl-L-lysine, N-acetylleucine, N6,N6,N6-trimethyl-L-lysine, and isovalerylglycine, were significantly enriched. These compounds are derivatives of CRC-associated parent amino acids such as tyrosine, valine, phenylalanine, lysine, leucine, and glycine, which were previously highlighted in Figures 3C, S7, and 6B. Their presence in the MS data is consistent with findings from NMR-based metabolomics (Figure 5B). Additionally, we detected N-α-acetyl-L-arginine, N-acetyl-arginine, and N(δ)- acetylornithine, which are metabolites involved in the arginine biosynthesis and degradation pathways in the co-culture supernatant (Figures 6). Arginine serves as a precursor for nitric oxide (NO) and polyamines; while polyamines promote tumor growth^92^, NO can exert either tumor-promoting or tumor-suppressive effects depending on its concentration^93^.

**Figure 6.**
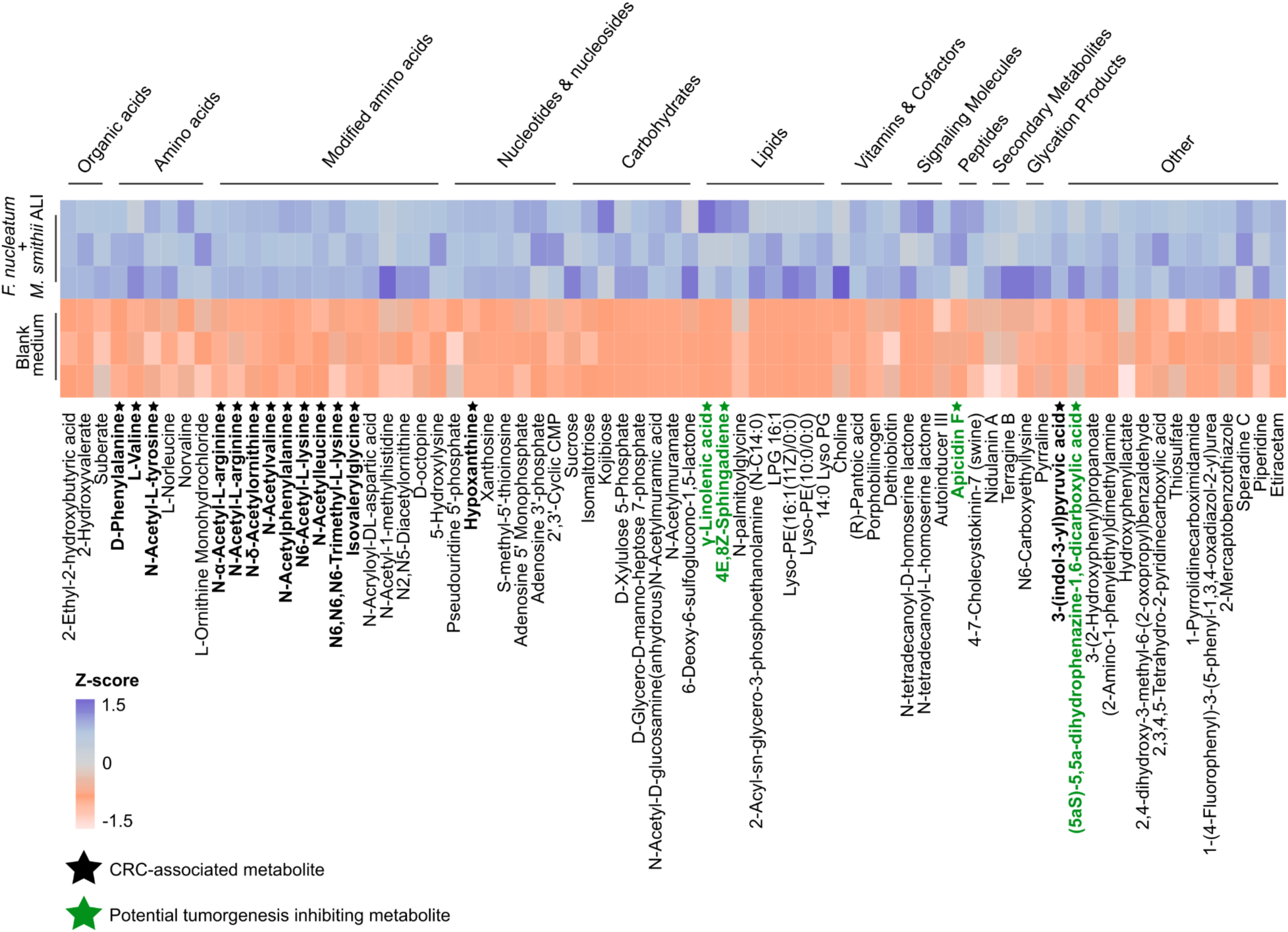
Mass spectrometry-based metabolomic profiling of the supernatant from *M. smithii* ALI + *F. nucleatum* co-culture. Heatmap showing significantly increased metabolites (FC > 1.5, t-test, *p*-adjusted < 0.05) detected by mass spectrometry in the co-culture of *F. nucleatum* and *M. smithii* ALI at t1 (24 h after addition of *F. nucleatum* to an established *M. smithii* ALI culture, coinciding with peak growth of both strains) compared to the blank medium (BHI+MS). Each cell represents the Z-score of the respective metabolite, calculated across all samples as the number of standard deviations from the mean. Metabolite concentrations were normalized using PQN and log₁₀-transformed to adjust for cell count differences and dilution effects. Metabolites with reported CRC-related promoting effects are highlighted in black, while those with potential tumorigenesis inhibiting effects are highlighted in green.

**Figure S12.**
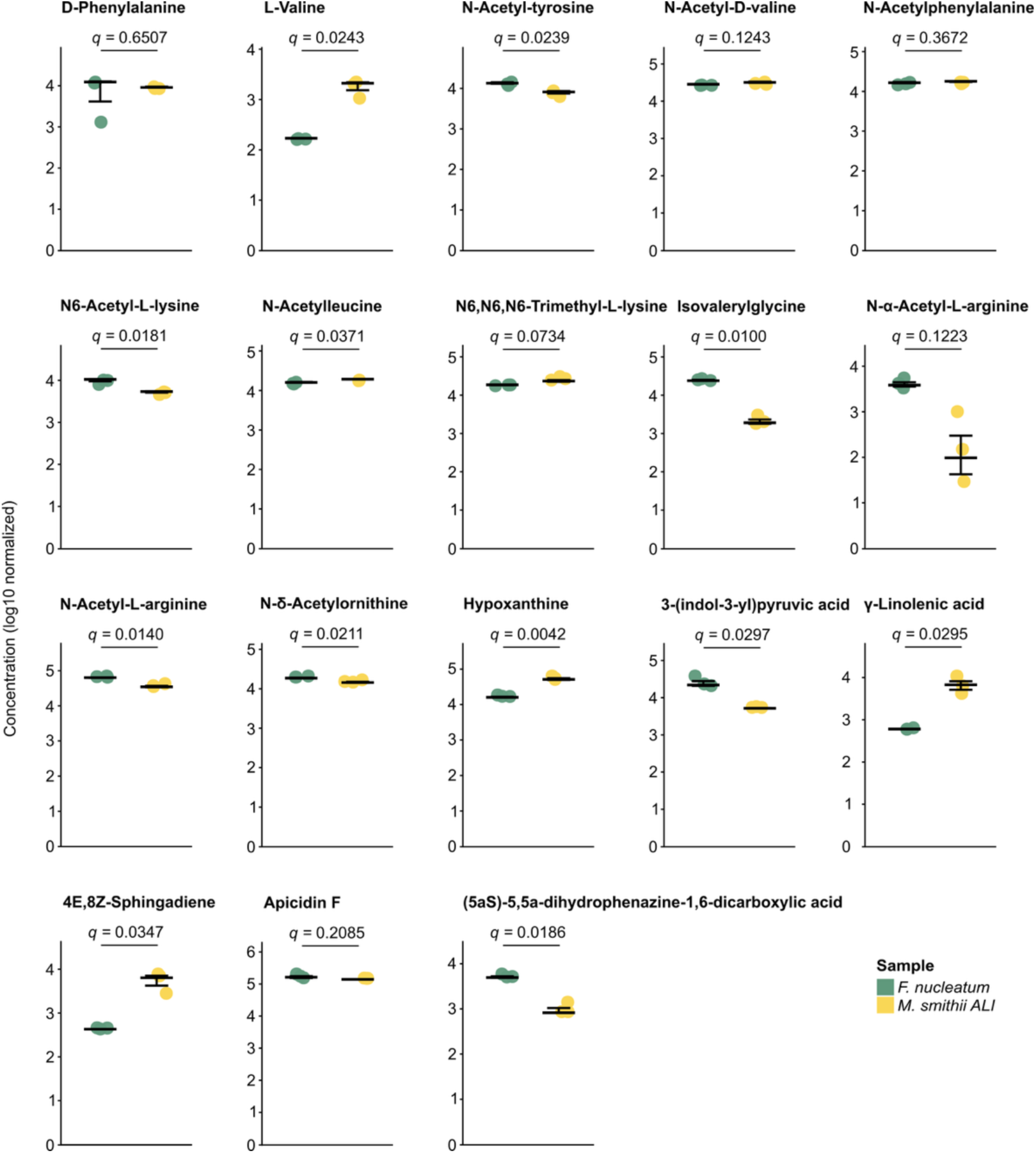
Scatter plot of metabolites with reported CRC-related effects (promoting or suppressive) based on literature. Metabolites were detected in the supernatants of optimally grown mono-cultures of *F. nucleatum* (BHI medium) and *M. smithii* ALI (BHI + MS medium). Metabolite concentrations were normalized using PQN and log₁₀-transformed to adjust for cell count differences and dilution effects. Statistical significance was assessed using a t-test, and *q*-values represent the FDR-adjusted *p*-values.

Hypoxanthine, a key intermediate in purine metabolism, was also detected in the co-culture supernatant of *M. smithii* ALI+*F. nucleatum*. This metabolite is widely produced and utilized by both host and microbial cells. Previous studies have associated microbial hypoxanthine production, particularly by *Peptostreptococcus stomatis,* another CRC-associated bacterium (Figure S7, Figure 3) with altered gut motility and serotonin release, processes that may contribute to CRC progression^87,94–96^.

Another interesting metabolite in the supernatant of the co-culture was 3(-indole-3-yl) pyruvic acid also known as indole-3-pyruvic acid (I3P). I3P supplementation has been shown to rescue tumor cells under tryptophan starvation, identifying it as a critical oncometabolite and therapeutic target^97^. Microbial production of I3P could circumvent tryptophan restriction, suggesting tumor– microbiome metabolic cross-talk that may promote cancer progression, particularly in gut-associated cancers like CRC, where gut barrier dysfunction could be present^98,99^.

Notably, several metabolites with known or potential antitumor activity were also identified in the co-culture supernatant (Figure 6). Among these, γ-linolenic acid, previously derived from *Lactobacillus plantarum* MM89, has been shown to induce ferroptosis in tumor cells, suggesting a novel therapeutic mechanism for targeting CRC^100^. Another detected metabolite, 4E,8Z-sphingadiene, belongs to the sphingadiene class, has demonstrated pro-apoptotic effects in colon cancer cells and has been shown to suppress intestinal tumorigenesis *in vivo*, highlighting its potential in cancer prevention^101^. Intriguingly, both γ-linolenic acid and 4E,8Z-sphingadiene were also significantly enriched in *M. smithii* ALI mono-cultures (compared to *F. nucleatum* mono-cultures), implicating the archaeon - and not the bacterial partner - as a direct source of potentially bioactive, antitumor metabolites (Figure S12, Table S11).

Apicidin F, a structural analog of the histone deacetylase inhibitor apicidin, was also present which has been previously isolated from the fungus *Fusarium fujikuroi^102^*. Its detection in the supernatant of co-culture suggests the potential ability of these two strains to also produce this compound (Figure S12). Notably, apicidin has shown anti-growth activity by inhibiting histone deacetylases in cancer cells^103^. Lastly, (5aS)-5,5a-dihydrophenazine-1,6-dicarboxylic acid, a derivative of phenazine-1,6-dicarboxylic acid, has been shown to be produced by strains such as *Pseudomonas aeruginosa*, and has demonstrated strong anticancer activity against various cell lines, including CRC cells like HT29^104,105^.

## Discussion

*Methanobrevibacter* species dominate the human gut archaeome. Their activity metabolizes H_2_ and CO_2_ into methane, thereby relieving lumen gas pressure, and sustains redox conditions that favor bacterial fermentation^14,35,106^. Although viewed as commensal, shifts in *Methanobrevibacter* abundance have been linked to various metabolic and gastrointestinal disorders, suggesting a potential role in host health^51,107^. However, most existing studies rely on low-resolution methodologies such as breath methane testing or 16S rRNA gene sequencing, which are limited to resolve archaeal taxonomy and to link archaeal species to physiological patterns^24^.

To address these limitations, we performed a meta-analysis of 1,882 shotgun-metagenomic datasets, drawn from 10 studies across 12 countries, and corrected for age, sex, and BMI. The datasets included different disorders, including colorectal cancer (CRC), type 2 diabetes (T2D), inflammatory bowel disease (IBD, including Crohn’s disease (CD) and ulcerative colitis (UC)), multiple sclerosis (MS), Alzheimer’s disease (AD), schizophrenia (SCZ), and Parkinson’s disease (PD).

Our analysis uncovered disease-specific alterations in gut archaeal communities. For instance, patients with CD showed a significant reduction in *Methanobrevibacter* spp., with the notable exception of Methanobrevibacter_A_sp900766745, which was significantly enriched in these patients. While the underlying mechanisms remain unclear, the reduced stool transit time in patients with CD may interfere with the slower growth rate of human gastrointestinal archaea. In contrast, SCZ patients exhibited a significant depletion of Methanobrevibacter_A_smithii, which confirms previously observed correlations of methanogen abundance and neurological function^3^.

Across cohorts, samples from CRC patients consistently demonstrated an increase in Methanobrevibacter_A_smithii and closely related taxa (e.g., Methanobrevibacter_A_smithii_A). While statistical significance varied between cohorts, the overall trend pointed toward the enrichment of methanogen in CRC, aligning with findings from a recent multicohort study^22^.

These findings hint to the limited but growing body of literature on the human archaeome in disease. Notably, only one of the investigated studies (Gupta et al.^64^) explicitly reported increased *Methanobrevibacter* abundance in CRC, while others (e.g., Feng et al.^57^) broadly implicated archaeal overgrowth without specifying the taxonomy. Our results were also in contrast with one of the included studies (those of Wallen et al.^69^), where they identified elevated *Methanobrevibacter* levels in PD patients. Our failure to replicate this finding, after correcting for sex, age, and BMI, underscores the importance of methodological consistency and the impact of metadata on archaeome and microbiome studies.

Variability in pipelines, reference genome databases, and cohort metadata contributes to inconsistencies across studies. These challenges highlight the need for consistent profiling frameworks and more comprehensive metadata collection, including antibiotic use, and medication history, which are often overlooked but likely influence archaeal dynamics. Dietary data were also not consistently reported across datasets, preventing us from accounting for potential differences in eating behaviors between case and control groups.

Methane, the primary end-product of methanogen metabolism, has been shown in animal models to slow intestinal transit^108^. Clinically, elevated baseline breath methane levels have been linked with constipation, which is a recognized risk factor for CRC^109,110^. These findings suggest a potential association between increased *M. smithii* abundance and CRC development. Notably, prior studies have reported enrichment of *M. smithii* in CRC patients even after considering gut transit time^22^, implying that factors beyond motility, including host–microbe interactions or microbial cross-talk, may play a role. However, methanogen abundance patterns remain susceptible to confounders such as constipation, which is common in CRC and can influence archaeal composition. The lack of standardized metadata, particularly regarding transit time, in existing cohorts limits our ability to fully resolve these effects and interpret associations^110^.

To investigate the functional contributions of *M. smithii*, we further explored its interactions with CRC-associated bacterial taxa. Given that *Methanobrevibacter* species rely on bacterial fermentation end products (H_2_ and CO_2_) for energy metabolism, their ecological function cannot be assessed individually. Therefore, we integrated archaeal and bacterial data, to examine how *M. smithii* may influence CRC-associated bacterial taxa and potentially contribute to colorectal carcinogenesis through microbe–microbe and potential host–microbe interactions.

While the loss of co-occurrence between commensal archaea and bacteria has been reported in CRC, *M. smithii* has shown positive co-occurrence associations with known CRC-associated taxa such as *F. nucleatum*, *B. fragilis* and *E. coli* in previous studies^22,35^. These bacteria are known to promote carcinogenesis by inducing DNA damage, inflammation, and activating oncogenic pathways, suggesting that CRC-enriched archaea and bacteria may synergistically contribute to CRC^111^. On the other hand, previous studies have reported significant co-exclusion patterns between CRC-enriched archaea and CRC-depleted bacteria, including butyrate-producing species such as *Clostridium beijerinckii* and *Clostridium kluyveri*, implying a potential antagonistic interaction in colorectal tumorigenesis^111^.

An interesting finding of our study based on metabolic modelling was the universal export of succinate by all CRC-associated bacterial partners and its predicted uptake by *M. smithii*, highlighting succinate as a key cross-fed metabolite. Additionally, *M. smithii* was predicted to export riboflavin in all archaeal-bacterial pairings and to import several CRC-associated amino acids, such as asparagine, glutamine, leucine, and arginine. These metabolic exchanges suggest potential cooperative networks that could influence both microbial ecology and host disease processes, beyond H_2_ exchange.

Experimental co-culture experiments validated our *in-silico* findings. *M. smithii* growth was significantly enhanced only in the presence of *B. fragilis*, while *F. nucleatum* and *E. coli* showed archaeon-induced early growth acceleration. In these conditions, the archaeon did not show a significant growth increase in return, indicating only mild commensal benefits or fully unidirectional support.

In all co-cultures, succinate levels were higher than in mono-cultures, supporting predictions from genome-scale metabolic models and confirming succinate as a shared cross-feeding metabolite. However, the overall changes in metabolism differed between bacterial partners. The *F. nucleatum–M. smithii* pairing showed the most extensive changes, with increased production of short-chain fatty acids (acetic, lactic, propionic, and butyric acids) and several amino acids (alanine, valine, isoleucine, leucine, phenylalanine, proline, and glycine), many of which have been linked to CRC^88,89,91,110^.

Untargeted mass spectrometry-based metabolomics of supernatants further confirmed these findings, showing not only elevated levels of these CRC-related amino acids but also their modified forms, such as N-acetylated derivatives (e.g., N-acetyl-tyrosine, N-acetylvaline, N6-acetyl-L-lysine).

Other detected metabolites are potentially directly connected to cancer-related pathways. For instance, arginine derivatives (N-α-acetyl-L-arginine, N-acetyl-arginine, and N-δ-acetylornithine) feed into nitric oxide and polyamine synthesis, both involved in CRC progression^91^. Interestingly, genes encoding cancer-related metabolites like polyamines have been shown to be more prevalent and diverse in gut and oral samples, affiliated with Euryarchaeota, especially methanogenic archaea^112^. Hypoxanthine, a purine compound known to affect gut function and tumor development, and I3P, an oncometabolite that supports tumor growth under tryptophan-starved conditions^87,94–96^, were also found in the supernatant.

The co-cultures also showed significantly high abundance of some compounds likely impairing tumorigenesis. These compounds included γ-Linolenic acid, which can trigger ferroptosis in tumor cells; 4E,8Z-sphingadiene, known to promote cancer cell death; apicidin F, a histone deacetylase inhibitor; and phenazine derivatives with strong anticancer activity^100,101,103–105^. Two of these compounds - γ-linolenic acid and 4E,8Z-sphingadiene - were significantly enriched in pure *M. smithii* cultures, indicating archaeal origin.

Our findings position *M. smithii* as an active metabolic participant in CRC-associated microbial networks. Through both cooperative and asymmetrical metabolic interactions, *M. smithii* modulates the abundance and function of key bacterial taxa and actively contributes to a chemically rich microenvironment that includes metabolites with tumor-promoting and tumor-suppressive potential. Understanding how host factors like immune status affect the balance of these interactions, including the archaea-metabolome will be key to determining their role in CRC development and therapy.

## Supplemental information

Document S1. Figures S1-S12

Document S2. Excel files containing additional data Tables S1-S11

## Resource availability

### Lead contact

Further inquiries and requests for materials or data should be directed to the lead contact, Christine Moissl-Eichinger.

### Materials availability

This study did not generate new, unique reagents.

### Data and code availability

Raw sequencing data for each published study can be accessed as described in Table 1. All generated and analyzed data (including outputs from Kraken2/Bracken, gapseq, PyCoMo, and results from ScyNet) are publicly available in our GitHub repository: https://github.com/CME-lab-research/archaeome-disease-profiling/. Raw metabolomics data as well as qPCR raw data are provided in Document S2. No original code was developed for this study.

## Acknowledgments

This research was funded in whole or in part by the Austrian Science Fund (FWF) [10.55776/P32697 (given to CME), 10.55776/CoE7 (CME and CD), 10.55776/F8300 (CME)]. C.T. reports a research grant by Bruker Switzerland AG.

For open access purposes, the author has applied a CC BY public copyright license to any author-accepted manuscript version arising from this submission.

We gratefully acknowledge the computational resources provided by the MedBioNode at the Medical University of Graz, funded by the Austrian Federal Ministry of Education, Science, and Research through the Hochschulrat-Struktur Mittel 2016 grant within BioTechMed Graz. We also thank the ZMF Core Facility Computational Bioanalytics team at the Medical University of Graz for their support. We thank Claire Lamb for assistance with the provision of the *Bacteroides fragilis* strain. R.M. was supported by the local PhD program MolMed.

## Author contributions

R.M. designed the study, collected data, performed bioinformatics, data analysis, plotting, and drafted the manuscript. A.M. co-designed the study. T.Z. and L.W. performed sample cultivation.

C.K. performed qPCR. K.F. assisted with plot preparation. P.M. performed *E. coli* isolation and confirmation. H.H. and T.M. performed NMR metabolomics. J.S. and C.T. performed MS metabolomics. D.P., K.H., and D.K. performed SEM. C.D. helped with data analysis. G.G and A.L. assisted with sample preparation. G.G., C.T., C.D., and A.L. commented on and revised the manuscript. C.M-E. designed and supervised the study, and drafted and revised the manuscript. All authors reviewed and approved the final manuscript.

## Declaration of generative AI and AI-assisted technologies in the writing process

Artificial intelligence (ChatGPT, OpenAI) was used solely to enhance the language and readability of the manuscript; no content was generated or interpreted by the tool.

